# Museum specimens reveal evolutionary drivers and functions of oral microbiome in mammals

**DOI:** 10.64898/2026.06.19.733412

**Authors:** Markella Moraitou, Piotr Rozwalak, Johnny Richards, Konstantina Saliari, Emmanuel Gilissen, Zena Timmons, Andrew C Kitchener, Olivier S. G. Pauwels, Richard Sabin, Phaedra Kokkini, Roberto Portela Miguez, Daniela Kalthoff, Fabian L. Kellner, Jaelle C. Brealey, Bas E. Dutilh, Michael D. Martin, Vebjørn Veiberg, Katerina Guschanski

## Abstract

The microbiomes of wild animals, despite being integral to host survival, remain largely unexplored, particularly beyond the gut. Here, using > 450 museum-preserved dental calculus samples from 34 ecologically and phylogenetically diverse mammalian species, most of them previously unstudied, we determine the main drivers of oral microbiome evolution. We show that, similar to the gut, the oral microbiome is shaped by host ecology, particularly diet, and to a lesser extent host phylogenetic relationships. It may contribute to host adaptation, by synthesising essential micronutrients and degrading potentially harmful compounds. We also find that some oral microorganisms consistently associated with mammals throughout over evolutionary time scales and provide evidence for a likely oral origin of some rumen taxa. Together our findings demonstrate the importance of this mostly overlooked microbial community for mammalian biology.

## Main

Microbiomes are essential for a number of host functions, including host digestion, development, immunity, behaviour, and ultimately fitness^1^. Throughout mammalian evolution, microbiomes have facilitated the colonisation of new ecological niches^1,2^, for instance by enabling the host to access previously inaccessible food sources such as plant fibre and carrion^3–5^.

Despite their functional importance, the factors shaping these communities are still poorly understood. The microbiome of the digestive system is distinct across mammal species and shaped by host diet, gut physiology and phylogeny^6,7^. The importance of these factors varies across host clades. For instance, primates show a strong effect of host phylogeny on their microbiome^8–12^, known as phylosymbiosis^13^, whereas bats do not^14^. Overall, the gut microbiome appears to be influenced by an interplay of evolutionary and ecological processes, including codiversification and coevolution, and environmental colonisation and host filtering, respectively^12,15–17^.

Host-associated microbiomes beyond the gut, such as the oral microbiome, remained largely unexplored, particularly in wild animals. The oral microbiome is a taxonomically and functionally complex community with a systemic impact on its host, best demonstrated by its role in several oral and non-oral diseases, including caries^18^, cancer^19,20^, Alzheimer’s^21,22^, heart infections^23^, and atherosclerosis^24^. Because of its role in nitrogen metabolism (reducing nitrate to nitrite), it is associated with vascular health^25^ and may even be linked to high-altitude adaptation in humans^26^ and non-human primates^27^. Studies on humans^28,29^ and wild animals^27,30–34^ suggest that, like the gut microbiome, the oral microbiome is shaped by diet and host phylogeny^35–37^ but is likely more exposed to environmental microbiota^38^.

Studying the oral microbiome of wild animals is hindered by the logistics and ethics of obtaining samples. As captivity alters the host microbiomes ^39,40^, captive animals are poor proxies for comparative studies. Dental calculus, the mineralised form of dental plaque^41,42^, found abundantly in museum specimens of a wide diversity of mammals^43,44^, provides a potential solution. Dental calculus forms through periodic mineralisation, which preserves microbiome DNA, allowing post-mortem sampling^45,46^. While widely used in anthropological and bioarchaeological research^32,46,47^, few non-primate studies have utilised this resource^31,34,44,48,49^. Here, using dental calculus, we generated and analysed metagenomes of 456 individuals from 34 species, containing representatives of most mammalian taxonomic orders. This unprecedented dataset allowed us to investigate the roles of host phylogeny, diet, digestive physiology and habitat on the taxonomic composition and functional capacity of the oral microbiome. It also provided a rare opportunity to assess the extent of host-oral microbiome codiversification, and to uncover ways in which oral microbiomes may contribute to host adaptation.

### Prey consumption is associated with higher microbiome diversity

Wild mammal oral microbiomes are underrepresented in public databases, so, to improve accuracy, we employed an assembly-based approach to taxonomic and functional profiling^50^. We relied on contigs ≥ 1000 bp in length instead of MAGs (metagenome assembled genomes) as historical DNA is highly fragmened^51^. The final dataset included 8,304 species-level taxa from 34 host species, representing eight taxonomic orders, four dietary classes (herbivores, frugivores, omnivores, and animalivores), and including three marine species (Table 1, Supplementary Tables 1 and 2, Supplementary Fig. 1d). Apart from domestic horses, sheep and pigs, all samples originated from wild individuals. Most microbial species belonged to the phyla Pseudomonadota, Bacillota and Actinomycetota, which are commonly found in human dental calculus^46^. Animalivores and omnivores showed a higher abundance of Bacteroidota and Synergistota than herbivores and frugivores (Fig. 1a).

**Fig. 1:**
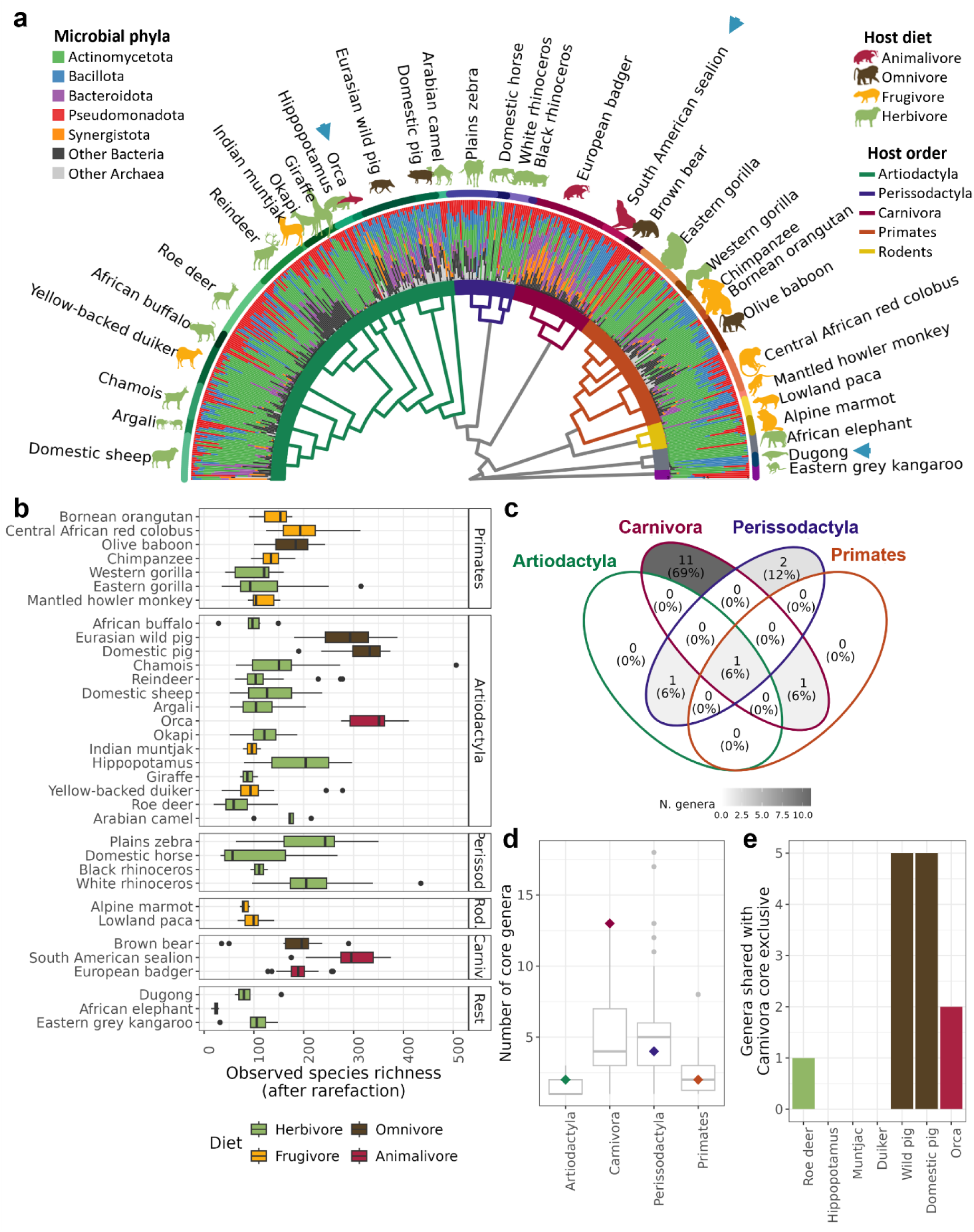
Taxonomic composition, richness and core of the mammalian oral microbiome. **a,** Microbiome composition plotted along the mammalian phylogenetic tree, with colours corresponding to microbial phyla. The branch colour reflects host taxonomic order. Animal symbols are coloured according to the dietary category and the three marine species are indicated by a blue arrow. Note that the phylogenetic tree did not include an entry for the domestic pig, so it is included under the same tree tip as its sister species, the Eurasian wild pig. **b,** Observed species richness per sample. Each boxplot represents a host species and is coloured by dietary category (red = animalivore, brown = omnivore, yellow = frugivore, green = herbivore). **c,** Venn diagram of the core genera in the four most densely sampled host orders: Artiodactyla, Carnivora, Perissodactyla and Primates. Only one microbial genus, *Actinomyces*, was found in the core of all four orders. **d,** Observed core microbiome size (number of genera), compared to the expected values given the number of host species per order. The diamonds indicate the original core microbiome size, whereas the boxplots show the results of 100 permutations of randomly sampling host species for each order, with the number of species fixed. The Carnivora core microbiome is much larger than expected by randomly sampling any three host species. **e,** Overlap between the exclusive Carnivora core microbiome (11 genera) and the core microbiome of seven artiodactyl species with different diets: herbivore roe deer (*Capreolus capreolus*) and hippopotamus (*Hippopotamus amphibius*), frugivore muntjak (*Muntiacus muntjac*) and yellow-backed duiker *(Cephalophus silvicultor*), omnivore European wild pig (*Sus scrofa*) and domestic pig (*Sus domesticus*), and the animalivore orca (*Orcinus orca*).

**Table 1.** Summary of the dataset, sorted by host order. Marine species are indicated with (m) in the ‘Common Name’ column. Orcas and false killer whales were considered as one species in all analyses.

| Common Name | Scientific Name | Host order | Diet category | Samples sequenced (retained) |
| --- | --- | --- | --- | --- |
| African buffalo | *Syncerus caffer* | Artiodactyla | Herbivore | 7 (7) |
| Arabian camel | *Camelus dromedarius* | Artiodactyla | Herbivore | 6 (6) |
| Argali | *Ovis ammon* | Artiodactyla | Herbivore | 10 (9) |
| Chamois | *Rupicapra rupicapra* | Artiodactyla | Herbivore | 18 (17) |
| Domestic pig | *Sus domesticus* | Artiodactyla | Omnivore | 12 (12) |
| Domestic sheep | *Ovis aries* | Artiodactyla | Herbivore | 29 (23) |
| Eurasian wild pig | *Sus scrofa* | Artiodactyla | Omnivore | 26 (26) |
| Giraffe | *Giraffa camelopardalis* | Artiodactyla | Herbivore | 8 (8) |
| Hippopotamus | *Hippopotamus amphibius* | Artiodactyla | Herbivore | 11 (10) |
| Indian muntjac | *Muntiacus muntjak* | Artiodactyla | Frugivore | 5 (4) |
| Okapi | *Okapia johnstoni* | Artiodactyla | Herbivore | 10 (10) |
| Orca (m) | *Orcinus orca* | Artiodactyla | Animalivore | 4 (4) |
| + False killer whale (m) | + *Pseudorca crassidens* |  |  | 1 (1) |
| Reindeer | *Rangifer tarandus* | Artiodactyla | Herbivore | 26 (25) |
| Roe deer | *Capreolus capreolus* | Artiodactyla | Herbivore | 35 (33) |
| Yellow-backed duiker | *Cephalophus silvicultor* | Artiodactyla | Frugivore | 23 (22) |
| Brown bear | *Ursus arctos* | Carnivora | Omnivore | 10 (10) |
| European badger | *Meles meles* | Carnivora | Animalivore | 38 (38) |
| South American sealion (m) | *Otaria flavescens* | Carnivora | Animalivore | 14 (13) |
| Eastern grey kangaroo | *Macropus giganteus* | Diprotodontia | Herbivore | 8 (7) |
| Black rhinoceros | *Diceros bicornis* | Perissodactyla | Herbivore | 2 (2) |
| Domestic horse | *Equus caballus* | Perissodactyla | Herbivore | 6 (6) |
| Plains zebra | *Equus quagga* | Perissodactyla | Herbivore | 24 (24) |
| White rhinoceros | *Ceratotherium simum* | Perissodactyla | Herbivore | 12 (12) |
| Bornean orangutan | *Pongo pygmaeus* | Primates | Frugivore | 3 (3) |
| Central African red colobus | *Piliocolobus foai* | Primates | Frugivore | 20 (20) |
| Chimpanzee | *Pan troglodytes* | Primates | Frugivore | 10 (10) |
| Eastern gorilla | *Gorilla beringei* | Primates | Herbivore | 28 (24) |
| Mantled howler monkey | *Alouatta palliata* | Primates | Frugivore | 5 (5) |
| Olive baboon | *Papio anubis* | Primates | Omnivore | 21 (21) |
| Western gorilla | *Gorilla gorilla* | Primates | Herbivore | 10 (10) |
| African elephant | *Loxodonta africana* | Proboscidea | Herbivore | 12 (11) |
| Alpine marmot | *Marmota marmota* | Rodentia | Frugivore | 11 (8) |
| Lowland paca | *Cuniculus paca* | Rodentia | Frugivore | 11 (10) |
| Dugong (m) | *Dugong dugon* | Sirenia | Herbivore | 5 (5) |
| **TOTAL** |  |  |  | **481 (456)** |

After rarefaction, most host species contained 100-200 species-level taxa on average (Fig. 1b), although animalivores and omnivores contained significantly more (ANOVA p-value < 0.05, not significant when accounting for host phylogeny; Supplementary Table 3, Fig 1b). Frugivory, habitat or ruminant digestion did not affect microbiome species richness, and neither did absolute latitude (Supplementary Table 3, Supplementary Fig. 2), despite it being associated with environmental microbiome diversity^52^ and biodiversity at large^53^. Faith’s phylogenetic diversity was strongly correlated with microbiome species richness and showed the same patterns (Supplementary Fig. 3).

To assess the conservation of the oral microbiome across mammals, we identified the core microbiome of each host species as the set of taxa with > 75% prevalence at > 0.1% relative abundance. The size of the core microbiome ranged from a single core genus in the herbivorous Western gorilla, African elephant, and domestic horse to 26 and 38 genera in the carnivorous badger and orca, respectively (on average 12 genera per host species; Supplementary Table 4). We also defined the core microbiome of each host order with at least three sampled species (Carnivora, Perissodactyla, Artiodactyla, and Primates) as the microbial genera that were core in ≥ 2/3 of host species. Only *Actinomyces* was conserved across all four host orders, with two additional genera (*Streptococcus* and *Arachnia*) being widespread across the dataset (present in more than a single host order’s core microbiome) (Supplementary Fig. 4). Therefore, these three genera are likely essential for oral biofilm in mammals.

Consistent with patterns of oral microbiome diversity (Fig. 1b), Carnivora had the most diverse core microbiome (13 genera from 5 phyla, only two shared with other orders; Fig. 1c, d, Supplementary Table 4). As three artiodactyl host species, all animalivores or omnivores (Supplementary Fig. 1a,b,c), were outliers in terms of diversity (Fig. 1b), we compared their core microbiomes to that of Carnivora, to understand if this higher diversity is driven by the same taxa. Indeed, these three host species shared more core genera with the Carnivora than their herbivorous or frugivorous artiodactyl relatives (Fig. 1e). However, the wild and domestic pigs shared five genera, whereas the orca shared only two (*Tannerella* and Aminobacteriaceae CAJPSE01) (Fig. 1e, Supplementary Fig. 4), suggesting the environmental acquisition of microbiota may also play a role. Owing to their marine habitat, the orcas may have relied on a different environmental reservoir to assemble the diverse microbiome typical of animalivores.

### Host diet is the primary driver of the oral microbiome, alongside habitat, digestive physiology and taxonomy

Redundancy analyses (RDA) found that host diet, digestive physiology, habitat and taxonomic order all shape oral microbiome composition but each to a different degree. Diet emerged as a major factor, with the first RDA axis separating animalivores and omnivores from frugivores and herbivores (Fig. 2a). Ruminants had a distinct microbiome, occupying the distal end of the herbivore cluster (Fig. 2a) and separating from herbivorous pseudoruminant Artiodactyla (Supplementary Fig. 5). Animalivores were associated with the phyla Bacteroidota and Synergistota and the genus *Flexilinea* (phylum Chloroflexota, previously reported as Anaerolineaceae bacterium oral taxon 439^54^). Ruminants were characterised by rumen- and ruminant-associated *Actinomyces* and *Propionibacterium*^55–58^ (Fig. 2b), consistent with previous studies of dental calculus^31,34^. Primates diverged along the second RDA axis (Fig. 2a) and, while being characterised by oral taxa (Fig. 2b) typical of the human mouth^59^, they still followed the same dietary cline along the first axis seem in the entire dataset (omnivores to herbivores, Fig. 2a), highlighting once more the importance of diet. The marine lifestyle also affected microbiome composition but less than host diet and taxonomic order (Supplementary Fig. 6).

**Fig. 2:**
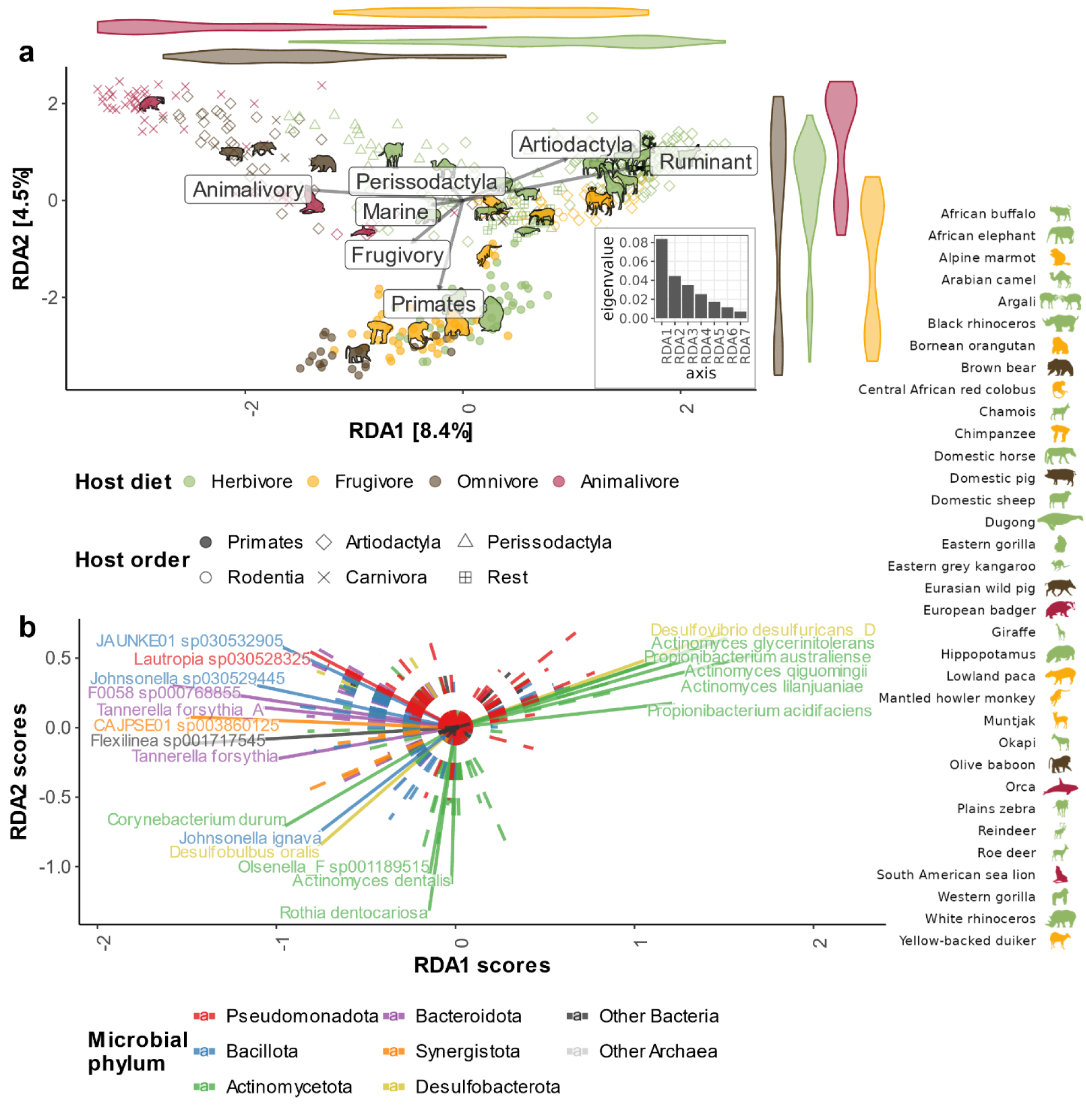
Redundancy analysis on CLR-transformed taxon abundances. **a,** Redundancy analysis on CLR-normalised abundances of species-level taxa for the entire dataset. Points represent samples and are coloured by dietary category. The length of arrows indicates the effect size of the explanatory variables reflecting host diet, taxonomy (the four most represented orders), marine lifestyle and ruminant digestion. The violin plots along the axes show the distribution of different dietary categories. The animal icons represent the centroids of each species (see legend on the right). The inset is a scree plot, showing the variance explained per axis. **b,** Loadings of 20 microbial species with the strongest correlation to the RDA axes in panel a (solid line), coloured by microbial phylum. Loadings for other unlabelled taxa are represented with dashed lines.

To formally test which microbial genera drive these differences in microbiome composition, we performed differential abundance analyses using two independent methods, PGLMM^60^ and MCMCglmm^61^. Supporting the RDA analysis and previous research^31,34^, PGLMM showed that the oral microbiome of ruminants was enriched for rumen-associated taxa (*Succiniclasticum*, *Porcinicola,* Erysipelotrichaceae RUG11795 and Methanomethylophilaceae RumEn.M2; Supplementary Table 5). A fourth genus, *Planktophila,* identified by both methods, is freshwater-associated^62^, so its enrichment in ruminants is not presently clear. Animalivory and frugivory were associated with more typical oral genera. Animalivores were enriched in *Flexilinea*, *Filifactor*, *Eubacterium*, and several uncultured taxa of the classes Clostridia and Saccharimonadia. Frugivores were only enriched in two taxa, *Pseudoramibacter* and the archaeon *Methanomethylophilus*. Finally, three genera were enriched in marine mammals: *Anaerococcus*, *Tractidigestivibacter* and *Enterococcus_E* (according to both methods; Supplementary Table 5). These are host-associated but not necessarily oral, possibly reflecting the poor characterisation of the oral microbiomes of marine animals. In addition, MCMCglmm identified six more marine-associated genera, including the porpoise-associated *Phocoenobacter*^63^ and *Psychrobacter*, typical of seawater^64^ and marine mammal skin^65^.

### Factors shaping the oral microbiome vary across mammalian clades

Multiple regression on matrices (MRM) with Aitchison’s distances confirmed that diet is a significant predictor of microbiome composition (Fig. 3a). However, host species identity emerged as the strongest predictor, especially for lower microbial taxonomic levels (e.g., species, genus; Fig. 3a, b). In comparison, host phylogeny had a considerably smaller effect and, like habitat and digestion, was only significant at intermediate microbial taxonomic resolutions (e.g., family, order; Fig. 3a, Supplementary Table 6). These patterns also held for other distance metrics, although Jaccard distances estimated a stronger effect of host phylogeny and ruminant digestion and PhILR distances estimated a stronger effect of habitat (Supplementary Fig. 7, Supplementary Table 6).

**Fig. 3:**
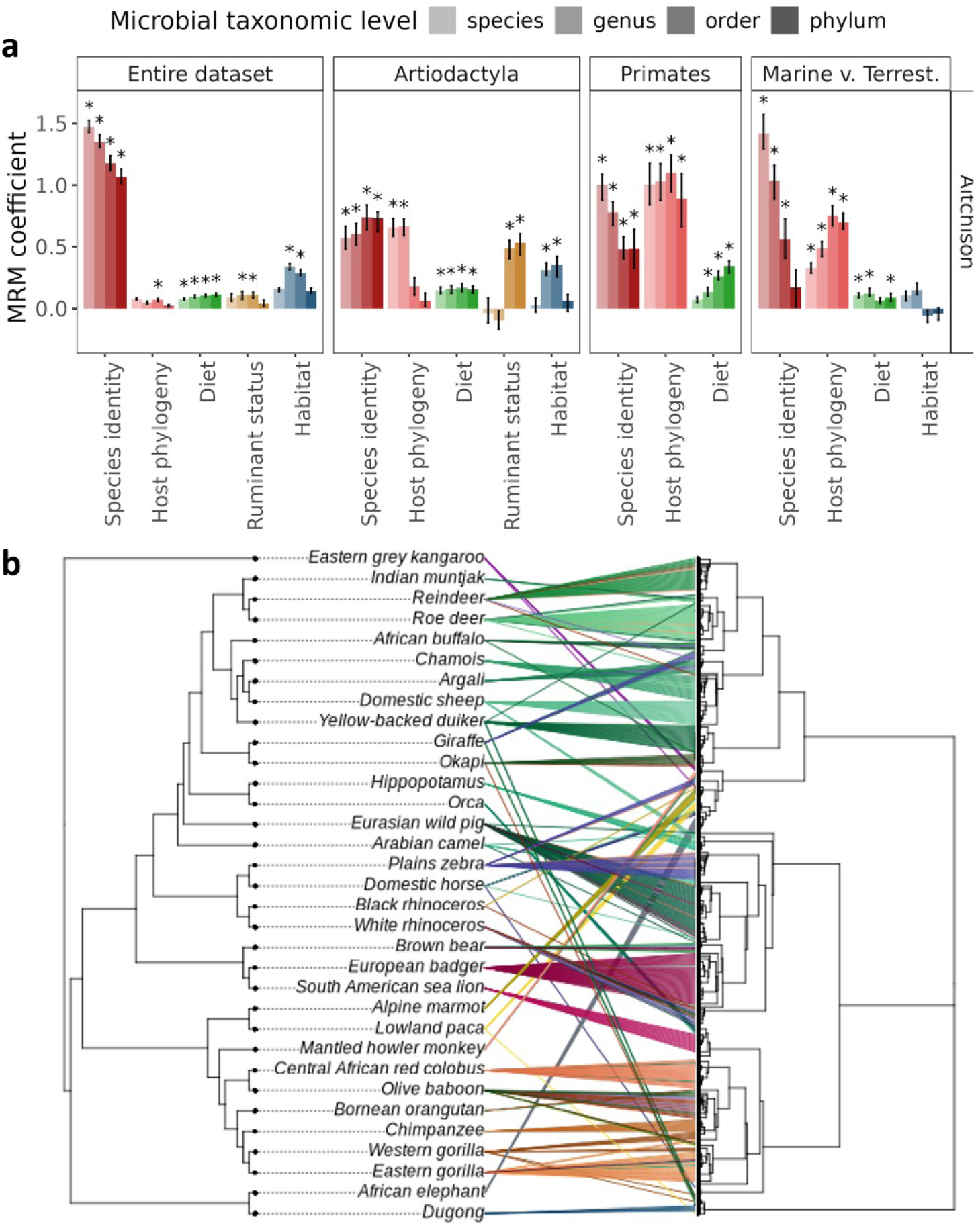
Host species identity is the main predictor of oral microbiome composition. **a,** Multiple regression on matrices using Aitchison distances performed for taxonomic resolutions at microbial species-, genus-, order- and phylum-level (for more levels of taxonomic resolution see Supplementary Table 6). The analysis was run on the entire dataset and on a subset of Artiodactyla, Primates, and marine mammals with their closest terrestrial relatives in the dataset (sealion—badger, dugong—elephant, orca— hippo/wild pig). Habitat signifies groupings into terrestrial or marine species. The bar heights represent the median coefficient and the error bars reflect the interquartile distance of 100 iterations, randomly subsampling five samples per host species. The asterisks reflect a median p-value < 0.05. **b,** Co-phylogenetic plot showing the congruence between host phylogeny (left) and microbial communities, represented by a neighbour-joining tree based on Aitchison distances (right). As the host phylogeny did not include an entry for the domestic pig, it is included under the same tree tip as its sister species, the Eurasian wild pig.

These relationships changed when Artiodactyla, Primates and marine mammals were considered separately (Fig. 3a, Supplementary Fig. 7). Most notably, the effect of host phylogeny was much more prominent, especially in Primates while diet remained significant but with a comparatively smaller effect size (Fig. 3a, Supplementary Table 6, Supplementary Fig. 7). Ruminant digestion showed increased importance in Artiodactyla in comparison to the entire dataset but, surprisingly, habitat (terrestrial or marine) appeared nonsignificant in the marine-terrestrial contrast, possibly because of variance absorbed by host species identity (Supplementary Fig. 8).

### Oral microbiome gene content hints at a potential role in host adaptation

Using the same set of contigs as for taxonomic profiling, we produced 23,679 unique gene annotations (after filtering) belonging to 99 KEGG pathways and 153 distillR functional traits^66^. Regardless of the metric used (gene abundances, pathway abundances or functional trait completeness), functional capacity profiles were congruent with taxonomic profiles, with the strongest congruence when comparing the abundances of microbial genera and genes (Procrustes p < 0.001; Supplementary Fig. 9). Host diet structured the functional potential of the oral microbiome, with animalivores and ruminants occupying the extreme ends of the RDA ordinations along the first axis (Fig. 4a, Supplementary Fig. 10). Again, MRM showed host species identity as the most important predictor of microbial functional potential, with host phylogeny and diet also playing a significant role (Supplementary Fig. 11).

**Fig. 4:**
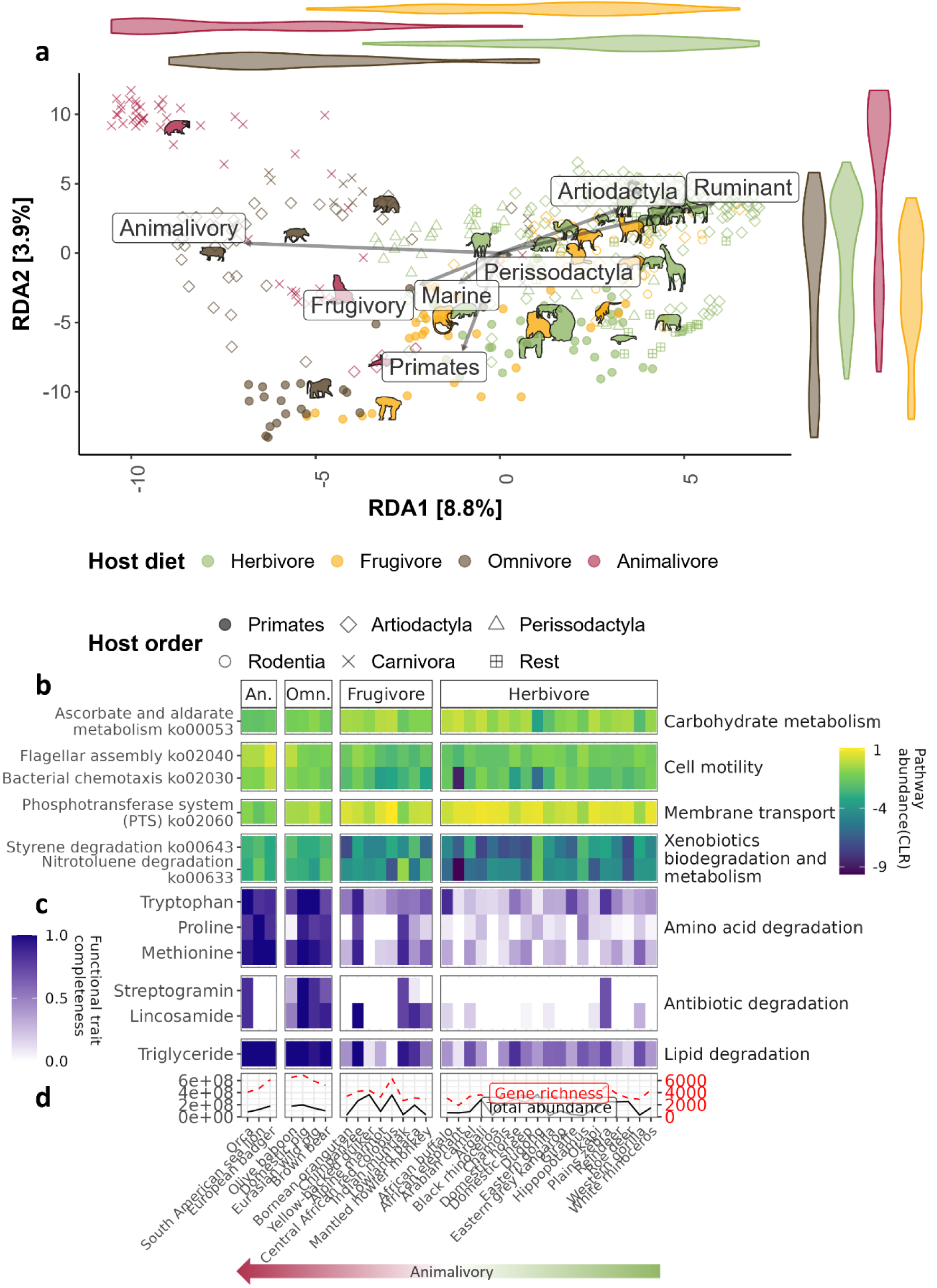
Diet drives functional differences in the oral microbiome. **a,** Redundancy analysis on CLR-normalised gene abundances. Points represent samples, with colour signifying dietary category and shape signifying host taxonomic order. The arrows indicate the effect of the constraints used for the analysis, specifically continuous variables describing diet (animalivory, frugivory), marine habitat, ruminant digestion and the most represented taxonomic orders (Artiodactyla, Perissodactyla, Primates). Violin plots alongside the axes show the distribution of different dietary categories. **b,** Mean abundances of the six KEGG pathways with the largest loadings along the first RDA axis grouped by pathway class (for sample plot see Supplementary Fig. 10a). **c,** Mean completeness of the six genome-inferred functional traits (calculated using distillR^66^) with the loadings along the first RDA axis grouped by function category (for sample plot see Supplementary Fig. 10b). **d,** Average total gene abundance (black solid line) and average gene richness (number of unique genes per sample, red dashed line). In panels b, c, d, host species are ordered left to right by decreasing proportion of animal content in diet and grouped into animalivores, omnivores, frugivores and herbivores.

Our findings suggest that the oral microbiome is adapted to host diet and may even influence host biology. Notably, the oral microbiomes of animalivores and omnivores had a higher gene diversity (Fig. 4d), in line with their taxonomic diversity (Fig. 1b), and appeared capable of utilising the nutrients most abundant in their environment, such as amino acids (specifically tryptophan, proline and methionine) and lipids (triglycerides) (Fig. 4c). These functions are likely to have an impact on host biology, as degradation of triglycerides by gut microbiota can reduce the risk of hypertriglyceridemia^67^, whereas degradation of methionine produces bioactive compounds relevant for the immune and neuroendocrine function of the host^68^. In addition, animalivore and omnivore oral microbiomes were enriched for degradation of xenobiotics, mainly acrylamide (part of the styrene pathway) and nitrotoluene (Fig. 4b, Supplementary Fig. 12a, b). Both compounds are of ecotoxicological concern^69,70^ and hosts at higher trophic levels (such as animalivores) are likely more exposed to them due to bioaccumulation. Finally, animalivore microbiomes contained more genes for flagellar assembly (Fig. 4b). This could be due to proposed differences in dental calculus biofilm composition^3^, although a similar finding in the gut microbiome was attributed to higher microbial motility due to intermittent feeding of the host^3^. These differences in community-wide functional capacity appear to be driven by differentially abundant taxa (Supplementary Table 5). For instance, the animalivory-associated *Flexilinea, Filifactor* and *Eubacterium* MAGs were enriched for xenobiotic degradation, flagellar assembly and lipid metabolism, respectively (Supplementary Fig. 13).

In herbivores functional profiles reflected the higher abundance of dietary soluble carbohydrates. The oral microbiomes of these animals carried more transporters for sugars and their derivatives from the phosphotransferase system (Fig. 4b, Supplementary Fig. 12d) as is typical of both the oral and gut microbiome under high sugar consumption^71,72^. The oral communities also had the capacity to utilise oxidised sugars via the glucarate/galactarate bacterial pathway (part of ascorbate and aldarate metabolism; Fig. 4b, Supplementary Fig. 12c). Consistent with this, differential abundance analysis associated herbivores and frugivores with the fructose and mannose metabolism pathway (animalivory coefficient < 0; Supplementary Table 7), particularly the metabolism of mannitol (Supplementary Fig. 12e), a common plant sugar linked to health benefits in ruminants^73^. Finally, enrichment analysis on MAGs found that the ruminant-associated *Succiniclasticum* and the frugivore-associated *Methanomethylophilus* were functionally enriched for porphyrin metabolism (Supplementary Fig. 13), particularly the branch for the biosynthesis of vitamin B12. This function is known from the herbivore gut and rumen microbiomes^74^ but our results suggest that the oral microbiome is also participating in the production of this essential for the host micronutrient.

Finally, differential abundance analysis identified six KEGG pathways with higher abundance in marine mammals, including thiamine metabolism (Supplementary Table 7). Thiamine production, which in our dataset appears enriched in the marine-associated *Anaerococcus* (Supplementary Fig. 13), is likely crucial for marine predators. While this essential micronutrient is abundant in animal tissues, an enzyme found in many fish species breaks it down before it can be absorbed, meaning that piscivores are at risk of thiamine deficiency^75^. This suggests that oral microorganisms may have facilitated the evolutionary adaptation of marine predators.

Nitrate reduction by the oral microbiome is relevant for cardiovascular health^25^ and has been proposed to be important for the high-altitude adaptation in gorillas^27^. To determine the generality of the phenomenon, we compared the abundance of nitrate reductases in three sets of high- and low-altitude species: gorillas (mountain gorillas — Western gorillas), rodents (Alpine marmots — lowland paca) and Caprinae (argali / chamois — domestic sheep). We also compared giraffes (*Giraffa camelopardalis*), whose long neck forces them to maintain chronic hypertension to keep blood flow to their brain^76^, against their sister species, the okapi (*Okapia johnstoni*). All high-altitude taxa were enriched in at least one nitrate reductase gene, but the global comparison (considering all genes together) was significant only with gorillas and Caprinae (Supplementary Fig. 14). No effect was seen in giraffes. Instead, some nitrate reductases were more abundant in the okapi.

Our analyses highlight the functional potential of the oral microbiomes, as characterised from fossilised dental calculus communities. Experimental work will be needed to fully establish their functionality.

### Widespread evidence for host-microbe codiversification in the oral microbiome

To assess if oral microbiota codiversify with the mammalian host, we relied on 575 dereplicated MAGs, including 525 high-quality (completeness ≥ 90% and contamination ≤ 5%) Bacteria and eleven Archaea (Supplementary Fig. 15, 16, Supplementary Tables 8, 9). We also included 38 Patescibacteria (or Candidate Phyla Radiation) regardless of completeness, as the reduced genomes of Patescibacteria^77^ may have led to an underestimation of genome completeness. Only 114 MAGs (19.8%) were classified at the species level, suggesting the presence of several novel species, and most were host species-specific (75%, N = 403 of 536), with an additional 12.5% (N = 67) being specific to host genus or host order (Supplementary Table 10).

Host-microbiome codiversification was widespread in the oral microbiome, affecting 17 of 59 tested MAG clades (permutation test: adjusted p-value < 0.05; Fig. 5a, Supplementary Table 11; N = 35 before adjusting for multiple testing) and spanned six phyla, including the four most abundant phyla in the mammalian oral microbiome: Actinomycetota, Pseudomonadota, Bacillota and Bacteroidota, as well as Desulfobacterota and Patescibacteria. Codiversification was more common amongst known oral (51%, 138 of 270 MAGs) and rumen taxa (45%, 14 of 31; Fig. 5b) than in gut- and soil-associated taxa (27% for gut and 45% for soil, although all codiversifying soil taxa were also reported in the oral microbiome), suggesting that the latter are likely transient.

**Fig. 5.**
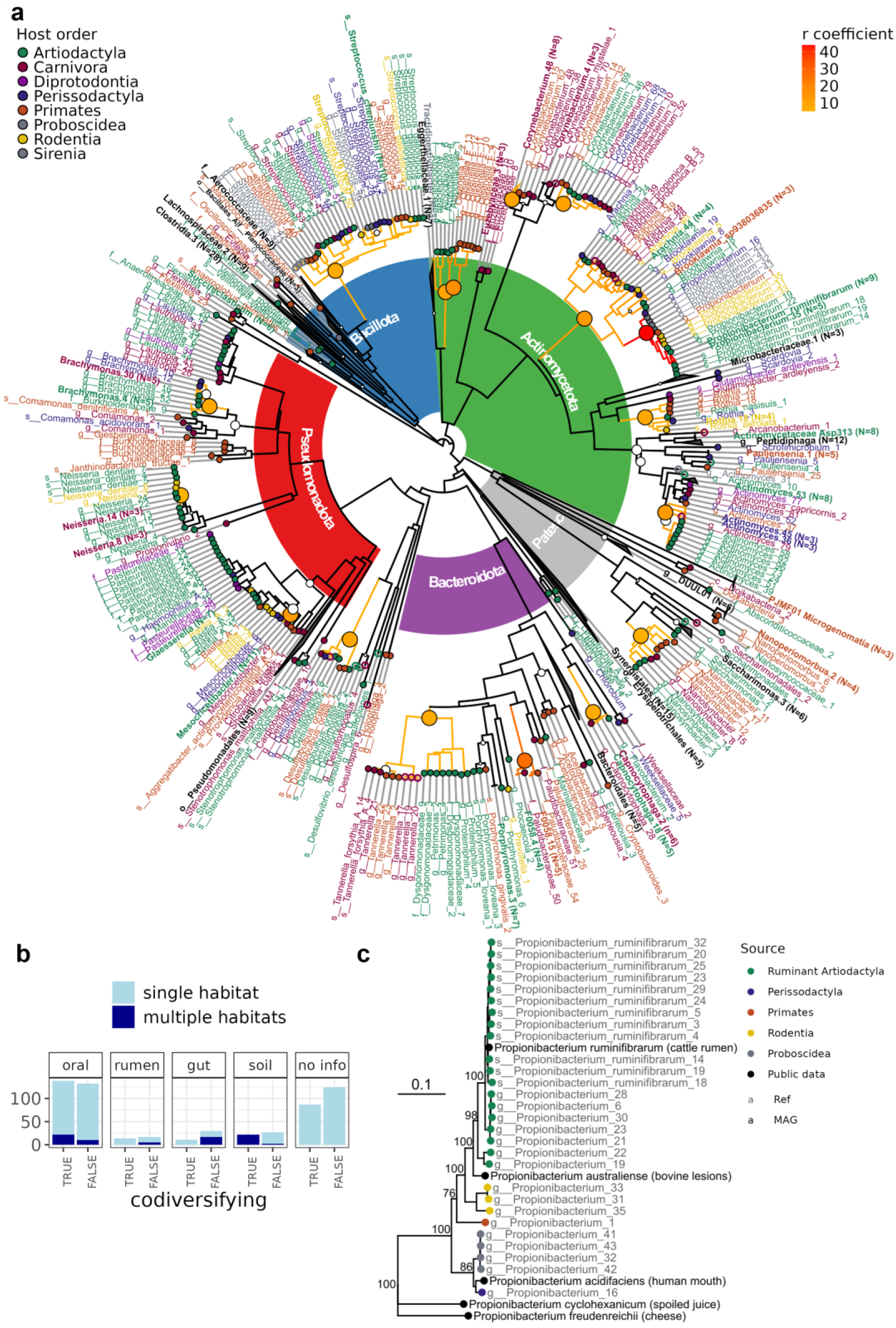
Host microbiome codiversification in the mammalian oral microbiome. **a,** Phylogenetic tree of 563 bacterial MAGs (525 high-quality MAGs and 38 members of Patescibacteria). For visualisation purposes, some non-codiversifying clades and clades with very short branches have been collapsed (labels in bold). From inside outwards: The five most represented phyla are coloured: Actinomycetota (green), Bacillota (blue), Pseudomonadota (red), Bacteroidota (purple) and Patescibacteria (grey). Tree clades and node circles are coloured by the value of the correlation coefficient r between host and MAG phylogenetic distances, with node circle size corresponding to log10-transformed adjusted p-values (only significant values are shown). The tree tips and labels are coloured according to the host taxonomic order (black for collapsed clades from several orders). Open circles represent marine hosts. **b,** Number of MAGs from codiversifying and non-codiversifying clades across habitats as reported in the Omnicrobe database, only retaining habitats that were mentioned in at least 30% of reports for that taxon. Taxa that have more than one habitat association are coloured in dark blue. **c,** Phylogeny of the genus *Propionibacterium*, including five references genomes (grey labels, with original isolation sources shown in brackets), suggesting potential oral origin of *P. ruminifibrarum*. Tip points for MAGs are coloured according to host order. Bootstrap support values are based on 1000 replicates. *P. ruminifibrarum* MAGs assembled from ruminant dental calculus are nested within unclassified *Propiopionibacterium* MAGs assembled from both ruminants and nonruminants. Some non-ruminant MAGs are closely related to *P. acidifaciens*, originally isolated from the human mouth. *Propionibacterium* reference genomes that are neither oral nor ruminant-associated appear as sister to this clade.

The three most common genera in the core oral microbiome of mammals, *Actinomyces*, *Streptococcus* and *Arachnia*, were found to codiversify (Fig. 5a), reinforcing the evidence for a strong host-microbe association. We observed the same for the oral genera *Rothia*, *Neisseria* and *Desulfobulbus* (Fig. 2, Supplementary Fig. 17a, b, c). Finally, the epiparasitic *Nanosynbacter* perfectly mirrored the host phylogeny, suggesting a strong dependence not only on the bacterial host, *Actinomyces* but also the mammalian superhost (Supplementary Fig. 17d).

Some codiversifying clades indicated that the oral microbiome may have served as a microbial source for the rumen microbiome. One such case is *Propionibacterium ruminifibrarum*, a bacterium isolated from the rumen^78^ but also found in reindeer dental calculus^34^. Several MAGs of this species, assembled from ruminant dental calculus, are nested among unclassified propionibacteria from both ruminant and non-ruminant hosts (Fig. 5c, Supplementary Fig. 17e). The phylogenetic placement of these clades suggests an origin in the mouth with a later translocation into the rumen. A similar pattern was observed within a clade of unclassified *Actinomyces*, a genus that is found in both the oral cavity and the rumen^57,59^ (Supplementary Fig. 17f). Here, a clade assembled from ruminants is nested within genomes assembled from Perissodactyla and Carnivora. Comparing the rate of molecular evolution between the ruminant and non-ruminant clades of these *Actinomyces* and *Propionibacterium* genomes (based on single-copy orthologues, N = 230 for *Actinomyces* and N = 378 for *Propionibacterium*), we found signatures of selection, suggesting adaptation to their ruminant hosts. In *Actinomyces*, several conserved genes (N=41) relevant for nutrient intake and energy production (e.g., membrane transport, starch and sucrose metabolism, oxidative phosphorylation and DNA metabolism) show stronger purifying selection in the ruminant clade than in the background branches (Supplementary Table 12). Conversely, the isoamylase gene, which is overall under positive selection (null dN/dS = 4.22), showed an even higher selection coefficient in ruminants (dN/dS = 6.89). In *Propionibacterium* two globally conserved genes showed weaker purifying selection in the ruminant clade, suggesting they may be undergoing genetic changes (adjusted p-value < 0.05; Supplementary Table 12).

### Bacteriophage families are shared among distantly related mammalian species

To characterise the viral component of the mammalian dental calculus microbiome, we identified 1,212 phage genomes representing 929 viral operational taxonomic units (vOTUs). Most vOTUs (n = 882) shared less than 95% average nucleotide identity with any of the 25.6 million sequences in viral sequence databases^79,80^, indicating that they represent largely novel species. Because most vOTUs were sample-specific, we grouped them at a higher taxonomic level using a gene-sharing network to cluster genomes into family-like groups (vFAMs). This analysis revealed that 19% of sequences belonged to 19 known vFAMs, 64% represented 277 previously undescribed vFAMs, and the remainder were classified as singletons (17%). Among the 25 vFAMs with more than ten genomes, only one was restricted to a single mammalian host (vFAM13 in badgers), whereas the remaining vFAMs were shared across multiple mammalian hosts, including distantly related species (Supplementary Fig. 18a). This pattern may reflect long-term associations between highly conserved bacterial lineages in dental calculus (Fig. 1a) and their viruses. Although phage families were broadly distributed and generally not associated with specific mammalian species, vOTUs often exhibited mammalian host species specificity (Supplementary Table 13), and certain phage lineages showed associations with animal diet (Supplementary Fig. 18b). Cataloguing this previously untapped viral diversity provides a resource for future exploration of bacteriophage evolution.

## Discussion

Here we present the largest comparative analysis of non-human oral microbiomes to date and demonstrate the suitability of museum-preserved dental calculus for comparative microbiome research. Consistent with studies of the gut microbiome^7,81^, diet shaped the taxonomic composition and functional capacity of the oral microbiome (Fig. 2, Fig. 4, Supplementary Fig. 10), reflecting its potential contributions to host biology and health. The animalivore microbiome, for instance, exhibited an increased capacity to degrade triglycerides (Fig. 4b), driven in part by an increase in *Eubacterium*, a taxon known for its anti-obesity effects^82^ (Supplementary Fig. 13). This microbiome-encoded function, alongside specific host genes^83–85^, may help protect animalivores from the negative health consequences typically observed in humans consuming a high-fat diet^86^. The animalivore oral microbiome was also enriched for the degradation of xenobiotics (Fig. 4b), similar to the gut, likely a response to contaminant bioaccumulation at higher trophic levels^87^. The high-protein diet of these animals seemed to drive an increase in genes encoding for degradation of amino acids (Fig. 4b), an activity linked to periodontitis in humans^88–90^. This is consistent with the higher abundance of periodontal pathogens (e.g., *Eubacterium*^91^, *Filifactor*^92^, Paludibacteraceae F0058^93^, *Tannerella*^94^) in the animalivore oral cavity identified both in our study (Fig. 2b, Supplementary Table 5) and previous work^31,49,95^. Their presence without signs of disease^44^ suggests that they are targeting dietary proteins rather than host gingival tissues and are therefore non-pathogenic in this context. These animalivory-associated functions, many of which appear adaptive, could explain why higher oral microbiome diversity emerged independently in carnivorans and artiodactyls that consume prey (Fig. 1b, d, e, Supplementary Fig. 4). This feature of the oral microbiome sets it apart from the gut, in which herbivores exhibit higher microbial diversity, possible owing to the demands of digesting plant fibre^81^.

In contrast, the herbivore oral microbiome appears adapted to an environment rich in soluble carbohydrates and was enriched for genes related to sugar metabolism and transport (Fig. 4). Two microbial taxa associated with frugivores and ruminants, respectively, were enriched for vitamin B12 (cobalamin) metabolism, an essential micronutrient that is absent in plants^74^. Similarly, oral communities of marine mammals were enriched in genes for vitamin B1 (thiamine) biosynthesis (Supplementary Table 7), another essential micronutrient often not accessible in piscivore diets^75^. Although these findings require experimental validation, which is not possible with dental calculus, they suggest a so far unrecognised contribution of the oral microbiome to the biosynthesis of micronutrients that cannot be obtained from diet, a functionality that has until now been primarily attributed to the gut and rumen microbiota^74^.

Beyond dietary adaptations the capacity of the oral microbiome to reduce nitrate has received attention for its potential cardiovascular benefits^25^ and putative role in high-altitude adaptation^27^. We found an increased abundance of nitrate reductase genes in mountain gorillas, confirming a previous study^27^, and furthermore extended this observation to chamois and argali, two high-altitude ungulates, and possibly to Alpine marmots (Supplementary Fig. 14). However, we found no evidence for increased nitrate reduction in giraffes, possibly because their hypertension tolerance is encoded in the host genome^96^.

Across the entire dataset host phylogeny was only a minor predictor of oral microbiome composition, consistent with findings in the gut microbiome^7,81^. This is best exemplified by the kangaroo, a marsupial that diverged from placental mammals ca. 160 million years ago. Yet, its oral microbiome is almost indistinguishable from that of other herbivores (Fig. 2a, 3b, 4a). Host phylogeny gained in importance when subsampling to specific host clades and especially in Primates (Supplementary Fig. 7, 8). This finding mirrors observations in the gut microbiome, which show strong phylosymbiotic signals in primates^8–10^ but not across the wider mammalian diversity^10,81,7^, suggesting that differences in relevant host traits (like diet, digestive physiology, etc.) can override the effect of phylogeny^14,97^.

Despite the weak phylogenetic signal overall, MAG-based analysis found several microbial clades that codiversified with mammals (Fig. 5, Supplementary Table 11), including several genera of the mammalian core oral microbiome, indicating strong host-microbiome associations over millions of years (Supplementary Table 18a). Some codiversifying bacteria are already known for their strong associations with the host: *Actinomyces, Streptococcus* and *Corynebacterium* are conserved in primates^47,98^ and *P. ruminifibrarum* has persisted in reindeer dental calculus spanning from the Pleistocene to the present day^34^. Our findings extend these host-microbe associations to an even deeper evolutionary time. This analysis also revealed likely oral origins of classical rumen taxa (*P. ruminifibrarum* and *Actinomyces*.6, Fig. 5c, Supplementary Fig. 17e, f). These ruminant bacteria show differences in the strength of selection compared to nonruminant clades (Supplementary Table 12), suggesting that they may be coevolving with the host instead of just passively codiversifying. Patterns of oral microbiome evolution also extended to the virome. While the same phage families were detected across mammals, their diversity reflected host diet, consistent with observations from the human and nonhuman virome^99,100^.

Overall, our findings reveal that the previously little explored oral microbiome may confer an array of important functions to the mammalian host, ranging from degrading excess lipids and xenobiotics, to synthesising essential micronutrients and providing physiological adaptations to extreme environments. We also find that certain oral microorganisms have been consistently associated with mammals over deep evolutionary time—codiversifying, and perhaps even coevolving with their hosts.

## Methods

### Samples

Our dataset comprised 482 dental calculus samples from 35 mammal species (1-38 samples per species, median = 11), 403 of which were newly sequenced and 80 were already published ^27,31,32,34,48,101^ (Supplementary Table 1). One of our newly sequenced samples was believed to be from an orca (*Orcinus orca*) specimen during sampling but was later confirmed to be a false killer whale (*Pseudorca crassidens*). Owing to the small sample size for the Cetacea (Delphinidae) clade and its unique ecology and phylogenetic position, this sample was combined with orcas for analyses (Table 1). Alongside dental calculus we also analysed 64 negative controls for decontamination purposes: ten environmental swabs from the museum collections, 27 extraction blanks (including three from Kellner et al^34^) and 27 library preparation blanks (see next section for more details).

### Data generation

Sampling and laboratory procedures were as described by Moraitou et al.^44^. In brief, dental calculus was sampled from museum-preserved specimens housed in five natural history collections: the Natural History Museum Vienna (Austria), which includes the Adametz collection of domestic animals, the Royal Museum for Central Africa in Tervuren (Belgium), the Natural History Museum London (United Kingdom), National Museums Scotland (United Kingdom) and the Royal Belgian Institute of Natural Sciences, Brussels (Belgium).

Special care was taken to minimise external and cross-contamination. Specifically, we wore disposable lab coats, face masks and two layers of gloves, with the upper layer changed for every new specimen. The sampling surface was decontaminated using 10% bleach solution and covered in a new sheet of aluminium foil for every new specimen. In every collection, we swabbed the collection environment to capture its microbiome, which could potentially contaminate the samples. These environmental swabs were processed, sequenced and analysed alongside the dental calculus samples.

Dental calculus was scraped off on a Whatman Weighing Paper with a sterile scalpel blade, aiming for ca. 50mg of material, and transferred to a 2 ml sterile microcentrifuge tube. To obtain sufficient material, multiple teeth were sampled per individual, often focusing on molars^44^. In rare cases, if the amount of dental calculus obtained from a single individual appeared insufficient, we pooled material from two conspecific individuals, matching their provenance, sex and age as closely as possible (Supplementary Table 1).

DNA was extracted and metagenomic libraries prepared in a specialised ancient DNA laboratory, following a strict decontamination regime. The DNA was extracted with the Dabney et al.^102^ protocol, with modifications by Brealey et al.^31^ and libraries were prepared according to a double-barcoding and double-indexing protocol ^103,104^. Sequencing was performed on twelve lanes of Illumina NovaSeq X Plus (PE 150 bp) (in two sequencing runs). Negative controls (environmental swabs, extraction blanks, library preparation blanks) were also sequenced and used to bioinformatically identify and remove contaminant taxa from our dataset.

### Metagenomic read pre-processing, assembly and binning

The newly sequenced metagenomic reads from this study were demultiplexed based on the inline barcodes using a custom script^31,104^ allowing one nucleotide error per barcode sequence. Then, for both newly sequenced and publicly available sequences, we used fastp^105^ to trim Illumina adapters, poly-G tails, unidentified and low-quality nucleotides (Phred score < 30, using sliding windows of 3bp), to clip 7bp-long barcodes on both ends of the read (only where present), and, finally, to merge the read pairs and retain only resulting reads ≥ 30bp. These reads were then deduplicated with dedupe.sh from BBMap^106^ to remove fully identical or fully contained reads, and mapped against a concatenated reference of the human genome, the PhiX virus genome, and the genome of the mammalian host (Supplementary Table 16) using bwa aln^107^. Negative controls (extraction blanks, library preparation blanks and environmental swabs) were mapped against the human and PhiX genomes only. The unmapped reads were extracted with SAMtools^108^. These metagenomic reads were evaluated with a source-tracking tool decOM^109^ to assess the oral microbiome proportion, using a custom source matrix including human oral, terrestrial mammal oral, marine mammal oral, rumen, skin, and sediment/soil microbiomes (Supplementary Information, Supplementary Table 14) and used to assemble contigs with MEGAHIT^110^ (default settings).

### Taxonomic and functional profiling of metagenomes

Contigs of length ≥ 1000 bp were first clustered at 98% nucleotide identity and 80% length overlap with the ‘easy-cluster’ function from MMseqs^111^ and then taxonomically and functionally profiled. Taxonomic profiling was based on the GTDB database^112^ after reformatting it for use with CAT^113^, using an approach adapted from https://github.com/MGXlab/gtdb2cat^114^. Genes were annotated with DRAM^115^, setting ‘--min_contig_size’ to 1000. To quantify contig abundances, we mapped the metagenomic reads using KMA^116^. We constructed taxon and gene abundance tables based on the number of mapped reads with at least 98% average identity, using contigs with at least 10% breadth of coverage for the taxon table and at least 50% for the gene table. We used KEGG (Kyoto Encyclopedia of Genes and Genomes), MEROPS (peptidase database) and CAZy (carbohydrate-active enzymes database) functional annotations. This contig-based approach allowed us to integrate the functional and taxonomic profiles, as we could directly link gene annotations to taxonomic assignments.

### Decontamination, filtering and normalisations of taxon and gene abundance tables

The abundance tables at species-level classification were analysed in R using the phyloseq package^117^, yielding 17,484 species-level taxa. Multiple sources of information were considered for decontamination. In summary, for each taxon, we considered relative abundance ratio in samples versus controls, previously reported habitats^118^, published lists of common contaminants^101,119^, and intensity of post-mortem DNA damage (detailed in Supplementary Information). Taxa with higher abundance in negative controls were also less prevalent in samples, more commonly reported in soil than in oral microbiomes and exhibited weaker damage patterns (Supplementary Fig. 19c, d, e, (Supplementary Table 15), which informed our thresholds for contaminant classification at relative abundance ratios ≤ 5 and prevalence ≤ 20% in every host species. In addition, we set to zero any abundance of a given taxon that accounted for < 0.01% of the total abundance in a sample. After this, 8,304 microbial species were retained for downstream analyses. Finally, samples with < 10,000 total reads mapped to taxonomically classified contigs, and/or an oral partition (human and animal oral partitions combined) smaller than the contaminant partition (soil/sediment and skin combined) were considered poorly preserved and removed from downstream analyses (Supplementary Fig. 20, Supplementary Table 1), which resulted in the only sample of Malayan tapir (*Acrocodia indica*) being excluded.

To account for compositionality^120^, abundances were normalised using CLR (centred log ratio, microbiome R package^121^) and PhILR (phylogenetic isometric log ratio^122^) transformations. The PhILR transformation was based on the GTDB bacterial phylogeny^54^ (release 220.0) and therefore excluded Archaea, to avoid the challenges of a shared Bacterial and Archaeal phylogeny.

For the gene table, contigs belonging to contaminant taxa (see above) were removed and the retained abundances summed per gene and sample. Then, genes with <100 reads across the dataset or ≤ 20% prevalence in every host species were removed. Samples identified as poorly preserved (see above) or containing less than 10^6^ functionally classified reads were also removed. The gene dataset was further summarised as KEGG metabolic pathways, using KEGGREST^123^, and as genome-inferred functional traits, using distillR^66^. We calculated pathway completeness as the proportion of genes present in the data, and abundance as the sum of the gene abundances, and excluded those with average completeness < 20% or not reflecting microbial metabolism (e.g., associated with human diseases, organismal systems or viruses). The distillR functional traits were calculated based on KEGG and Enzyme Commission annotations, and reflect metabolic capacities, such as the ability to degrade and biosynthesize different compounds (with values ranging from 0 to 1). They were hierarchically organised within functions and ultimately clustered into three domains: degradation, biosynthesis and structure.

### Host diet and phylogeny

To obtain quantitative estimates of mammalian diet, we consulted two sources. Where numerical variables were needed, we used dietary item proportions from the EltonTraits database^124^, which presents dietary composition in terms of ten dietary item types (e.g., invertebrates, fruit, seeds). We summed all dietary items indicating animalivory (e.g., invertebrates, endotherms, scavenge) into a single item. The diet of included study species was described as dietary proportions of animal, fruit, seeds and other plant material (Supplementary Fig. 1b, Supplementary Table 2). The first two proportions, referred to as animalivory and frugivory, were used in statistical tests (see next sections). When discrete dietary categories were needed, we used the diet clusters from the MammalBase database^125^, which were calculated based on estimated macronutrient content (e.g., crude protein, crude fibre, etc; Supplementary Fig. 1a, Supplementary Table 2), after adjusting them to reconcile them with EltonTraits and common knowledge about our study species. Specifically, based on ordination of dietary items proportions, the giraffe (*Giraffa camelopardalis*) was reclassified from frugivore to herbivore, the Eurasian wild pig (*Sus scrofa*) and domestic pig (*Sus domesticus*) from herbivore to omnivore, and the Alpine marmot from herbivore to frugivore (Supplementary Fig. 1c, Supplementary Table 2).

Host phylogeny was based on the consensus tree of 100 sampled subtrees (Mammals birth-death node-dated DNA-only trees) obtained from vertlife.org^126^. As the tree did not contain the domestic pig (*Sus domesticus*), for phylogenetic analyses they were treated the same as wild pigs (*Sus scrofa*).

### Statistical analysis of alpha diversity

To allow for valid comparisons of microbiome diversity, all samples were rarefied to 20,000 reads mapping to classified contigs. This threshold was chosen after plotting rarefaction curves using the vegan R package^127^ and identifying the number of classified reads at which the species richness reaches an asymptote. Then, we calculated species richness using phyloseq^117^ and Faith’s PD (phylogenetic diversity^128^) using the picante R package^129^.

We tested for the effects host diet (animalivory and frugivory), habitat (terrestrial vs marine), and ruminant digestion on microbiome species richness using a linear mixed-effects model, implemented in lme4^130^, with these host traits as fixed effects and host species as a random effect. Also, to account for host phylogeny, we implemented a similar model in MCMCglmm^61^ with weakly informative priors set to 0 (iterations = 40,000, burn-in = 20,000, thinning = 50), including, in addition, a phylogenetically structured variance/covariance matrix.

Finally, we assessed the presence of latitudinal diversity gradients using linear mixed models of species richness with either absolute values of latitude (estimated where sufficient geographical metadata were available) or a categorisation into tropical/non-tropical as predictors (Supplementary Table 2), and host species as a random effect.

### Statistical analysis of taxonomic and functional composition

Taxonomic and functional beta diversity (the later encompassing genes, pathways, and functional traits) was visualised using redundancy analyses (RDA) with the microViz^131^ and the rphylopic R packages^132^. Aitchison distances were used for taxon, gene and pathways abundances, and Euclidean distances were used for functional trait completeness. The constraints included host taxonomic order (for orders with at least three species: Artiodactyla, Primates, Perissodactyla, Carnivora), diet (degree of animalivory and frugivory), habitat (terrestrial or marine), and ruminant digestion. The fit of the model was tested using a step function and variance inflation factors. The analysis was repeated for subsets of the dataset, modifying the constraints accordingly (for instance, ruminant and pseudoruminant digestion were used in the Artiodactyla subset). The congruence of taxonomic (species- and genus-level abundances) and functional composition (pathway and gene abundances) were tested using the ‘procrustes’ and ‘protest’ functions of the vegan package.

The effects of host phylogeny and traits on taxonomic and functional composition were tested using multiple regression on distance matrices (MRM) with the ecodist R package^133^. All predictors were encoded as distances: host phylogeny, diet, habitat, ruminant digestion and host species identity (Supplementary Information). For taxonomic compositions, dissimilarities were calculated for various taxonomic resolutions from species to phylum (Supplementary Information), and using three distance metrics: Jaccard distances (reflecting differences in presence-absence), Aitchison distance (differences in abundances), and Euclidean distances on PhILR abundances (phylogeny-weighted differences in abundances) and were all centred and scaled. For pathway and gene composition, dissimilarities were calculated using Jaccard and Aitchison, and for functional trait completeness using Jaccard and Euclidean. To account for different sample sizes across species, this analysis was caried out by subsampling five samples per host species with replacement, for 100 iterations per model. It was also performed both on the entire dataset and in subsets, as described for the RDA.

### Differential abundance analysis

To identify differentially abundant taxa and pathways, we used two approaches: a) a two-pronged approach adapted from Youngblut et al.^12^, which tested separately for host traits accounting for phylogeny, and for phylogeny accounting for host traits, and b) a single model in MCMCglmm^61^ (also described by Sweeney et al.^134^). Differential abundance analyses were run on the top 200 most abundant genera (according to a weighted average of CLR-normalised abundances to mitigate biasing towards species with more samples) and for all pathways. First, for the two-pronged approach, we ran PGLMMs (phylogenetic generalised linear mixed models) using the phyr R package^60^ to test for the effect of host traits (degree of animalivory and frugivory), while accounting for phylogenetic relatedness by including host species as random effect with a variance/covariance based on phylogenetic distances. To test for phylogeny accounting for host ecological traits, we regressed out these same predictors using a linear model and then ran a Pagel’s lambda test using the ‘phylosig’ function from the phytools R package^135^. We adjusted p-values for multiple testing using the Holm method.

Second, we implemented a single model in MCMCglmm^61^ (40,000 iterations, 20,000 used as burn-in, 50 thinning interval), which, similarly, included host traits as fixed effects and host species as a random effect but in addition included phylogenetically-structured variance/covariance matrix. Using the MCMCglmm output we calculated phylogenetic lambdas (λ)^136^. PGLMM and MCMCglmm yielded generally consistent results for the correlation coefficients but not the values of lambda (Supplementary Fig. 23).

To link taxonomic differential abundances to changes in functional potential, we identified high-quality metagenome-assembled genomes (MAGs; see below for details) from the differentially abundant genera. When possible, we used MAGs assigned to these same genera, however for certain genera, no MAGs were assigned the same taxonomy, so we used MAGs using either family or order level classification. We then performed gene set enrichment analysis (GSEA) on these MAGs using the clusterProfiler R package^137^, with the background the list of all genes identified in the dataset.

### Core microbiome

We defined the core microbiome of a host species as the set of genera with at least 75% prevalence at 0.1% relative abundance, and the core microbiome of a host order (for those containing at least three species: Artiodactyla, Carnivora, Perissodactyla, Primates), as the set of microbial genera that are in the core microbiomes of two thirds of species in that order. As these orders varied in the number of species, we evaluated the robustness of these results by permuting the host species in each order, while maintaining the same order (100 iterations).

### Metagenome assembled genomes and codiversification analysis

Contigs were binned into metagenome-assembled genomes (MAGs) separately for each sample, using the nfcore/mag pipeline (v 3.2.1-gb582aae) with its ancient DNA module activated^138^ (and without GTDBtk, Prodigal, Prokka, Metaeuk and CONCOCT). This module assesses the presence of deamination damage patterns^138^ using PyDamage^51^ and corrects the nucleotide misincorporations by generating consensus sequences with Freebayes^139^ and BCFtools^140^. Through this pipeline, we assembled bins using MetaBAT2^108^ and MaxBin2^141^, and the resulting bins were refined with DASTool^142^ to keep only the highest quality nonredundant bins. In some cases, only one of the two binning tools successfully generated bins, in which case bin refinement was not performed and the output from the successful software was used in downstream analysis.

MAGs with ≥ 50% completeness and ≤ 10% contamination (as evaluated with ‘lineage_wf’ from CheckM^143^), were assigned taxonomy with the ‘classification_wf’ function from GTDBtk^112^ and used to construct de-novo phylogenies with the ‘denovo_wf’ function. MAG abundances were estimated by mapping the metagenomic reads to a combined reference of MAGs dereplicated at 99% identity (with ‘dereplicate’ from dRep^144^), using the KMA mapper^116^. Phylogenetic trees were visualised in R using the ggtree^145^ and ggtreeExtra^146^ packages.

Focusing on high-quality MAGs (completeness ≥ 90% and contamination ≤ 5% according to CheckM results), as well as members of Patescibacteria regardless of completeness (as they have reduced genomes and hence likely underestimated completeness), we tested for microbe-host codiversification. To do this, we first determined MAG presence in each host species by mapping the metagenomic reads from each sample against the collection of high-quality MAGs. When using samples from which the MAGs were assembled, we regularly recovered >10,000 mapped reads, breadth of coverage > 75%, and average nucleotide identity > 98%. Hence, we used these thresholds as the baseline to determine the presence of a MAG in a given sample (Supplementary Fig. 21).

Next, similar to Moeller et al.^147^, we scanned across the MAG phylogenetic trees (bacterial and archaeal) calculating the correlation of MAG and host phylogenetic distances (r correlation coefficient, one-sided Pearson test with alternative = “greater”) for all clades with depth equal or less than 30% of total tree depth (depth defined as median tip to node distance). Only MAG clades with at least five MAG genomes across three host species were tested. We then selected the node with the strongest correlation per lineage and estimated p-values by permuting the tips of the MAG clade 500 times (pseudocount of 10^−5^ added to prevent 0 values) (Supplementary Fig. 22). Clades showing codiversification were visualised using the ‘cophylo’ function from the phytools R package^135^. For selected clades, we validated the GTDBtk de novo phylogeny using IQ-TREE 2^148^ on the protein alignment already generated by GTDBtk with the WAG+F+I+R2 model and 1000 bootstrap replicates.

For some of the codiversifying clades, we tested for differences in the rate of molecular selection using a branch model implemented in CODEML from the PAML package^149^, adding also related reference genomes from GTDBtk. This required downloading and annotating them with DRAM. Then, single-copy orthologues were identified using OrthoFinder based on the amino acid gene sequences from DRAM. Orthogroups that contained more than one unique gene description were excluded (for instance, “3-isopropylmalate/(R)-2-methylmalate dehydratase small subunit [EC:4.2.1.33 4.2.1.35]” and “Aconitase C-terminal domain [PF00694.22]”). The nucleotide sequences of the retained orthogroups were aligned using MAFFT^150^ and the alignments filtered to contain only codons with no gaps, using a custom script from Diepeveen et al.^151^. Using the filtered alignments, we tested differences in the rate of molecular evolution using dN/dS tests implemented in CODEML, parallelized using GNU parallel^152^. Specifically, two models were run, a null model (model=0) assuming as single dN/dS ratio, as well as a branch model (model=2) assuming different dN/dS ratios between the foreground and background branches. The rest of the parameters were: seqtype = 1, ndata = 1, cleandata = 0, NSsites = 0, CodonFreq = 6, estFreq = 0, clock = 0. Likelihood ratio tests were used to compare the null models against the branch models.

### Bacteriophage identification

Bacteriophage sequences were identified using geNomad^153^ v1.11.2. Genome quality was estimated using CheckVhttps://paperpile.com/c/y7cNZn/2WUe^154^ v1.0.1 with database v1.5. Phage genomes predicted as medium-quality, high-quality, or complete were subsequently clustered into vOTUs at the species level (95% average nucleotide identity over 85% alignment fraction) and genus level (70% average nucleotide identity over 85% alignment fraction) using Vclust^155^, and into taxonomically described and novel family-level groups using gene-sharing network implemented in vContact3^156^ v3.2.0 with the RefSeq database v232. The novelty of vOTUs was assessed by clustering mammalian dental calculus phages with sequences from the Vire^80^ and metaVR^79^ databases at species-level thresholds. Putative bacterial hosts were predicted for phage genomes using iPHoP v1.4.2 with the Jun_2025_pub_rw database.

## Data Availability

Newly generated sequencing data is available at the European Nucleotide Archive under project accession number XXXX. Contig and MAG sequences have been uploaded to Zenodo XXXX. Sequence pre-processing scripts are available at https://github.com/markella-moraitou/mammal_oral_microbiome_preprocessing, scripts for statistics and figures at https://github.com/markella-moraitou/mammal_oral_microbiome_stats and phage analyses at https://github.com/rozwalak/mammalian_oral_phages.

## Acknowledgements

We would like to express our gratitude to the following individuals for their contributions to this project: Annie G. West, Dominik W. Melville and Emma Diepeveen for offering advice on different aspects of the methods; Arve Lee Willingham for drawing our attention to *Flexilinea* phylogenetic pattens; Frank E. Zachos, Alex Bibl, and Tim Langnitschke from the Natural History Museum in Vienna (Austria), and Dominique Fonck and Mathys Rotonda from the Royal Museum for Central Africa (Belgium) for welcoming us in the collections and assisting during sampling; Gunilla Engström and E-Jean Tan from Uppsala University (Sweden) for their support during the laboratory work; Yanara Marincevic-Zuniga and the rest of the team at the SNP&SEQ Technology Platform, Science for Life Laboratory, Uppsala (Sweden) for their expertise and support in sequencing (project codes WI-3800 and XC-3946).

