## Supplementary Information and Figures for "Museum specimens reveal evolutionary drivers and functions of oral microbiome in mammals"

### Supplementary Methods

#### decOM sources

We performed source tracking with decOM, a kmer-based microbial source tracking tool developed specifically for dental calculus metagenomes[1]. Because the default decOM source matrix only includes human oral metagenomes, we constructed a custom source matrix which also included oral metagenomes from wild mammals, as well as rumen metagenomes from wild and domesticated animals. More specifically, this source matrix consisted of 478 metagenomes:

- 177 human oral metagenomes from the default decOM source matrix, including both ancient (mostly dental calculus) and modern samples (mostly plaque, as well as saliva, gum and dental calculus),
- 49 terrestrial mammal oral metagenomes, specifically dental calculus from brown bears and reindeer[2,3],
- 52 marine mammal oral metagenomes, specifically gingival sulcus from dolphins and a harbour seal (49 metabarcoding and three shotgun metagenomic)[4,5],
- 40 rumen metagenomes from moose, sheep and cow[6–8],
- 79 sediment/soil metagenomes from the default decOM source matrix,
- 81 skin metagenomes from the default decOM source matrix.

#### In-silico decontamination

We considered multiple sources of information to identify and remove likely contaminants in our dataset. For every microbial taxon that was found in both dental calculus samples and controls (including environmental swabs and laboratory controls), we considered its relative abundance ratio in samples versus controls, its prevalence and average relative abundance in samples, the habitats it had previously been reported in, its presence in published lists of common contaminants[9,10] (note that some of the listed genera are also common in the oral microbiome e.g. *Streptococcus*, *Propionibacterium*, *Pseudomonas*), as well as the intensity of post-mortem DNA damage (Supplementary Table 15, Supplementary Fig. 19).

The sample-to-control abundance ratio was obtained by calculating the weighted average relative abundance of a taxon in dental calculus samples (first averaged within host species, and then across, to account for the different number of samples) and dividing it by the average relative abundance in negative controls (laboratory blanks and environmental swabs. Prevalence was calculated as the proportion of samples per host species that contained a given taxon at > 0.01% relative abundance (Supplementary Table 15, Supplementary Fig. 18).

The information on previously reported habitats was retrieved from the Omnicrobe database[11]. To match the taxa in our dataset with the Omnicrobe database, we first obtained NCBI taxonomic IDs using the taxize R package[12]. When we could not match the species name to the NCBI taxonomy database, we tried again with only the genus name. Then, using these IDs, we searched the Omnicrobe to retrieve the how many times each taxon was reported in the following pre-defined habitats: "laboratory equipment", "marine water", "deep sea", "soil", "mammalian", "wild animal", "mammalian livestock", "biofilm in natural environment", "host associated biofilm", “skin”, "mouth", "dental plaque", "gut" and "rumen".

The damage pattern estimates were extracted from the output of the PyDamage[13] module of nf-core/mag[14]. Specifically, we used the “damage_model_pmax” parameter, which indicates the maximum amount of damage and the 5’ end of the read. Because the output included damage_model_pmax estimates per contig, we summarised the values per microbial taxon by averaging.

Considering all these independent lines of evidence for decontamination, we found that taxa that were more abundant in negative controls tended to also have lower prevalence across samples (Supplementary Fig. 19c), were more often reported in soil than in oral microbiomes (Supplementary Fig. 19d), and showed less evidence for post-mortem DNA damage patterns (Supplementary Fig. 19e). Based on this information, we chose to retain only taxa that had an average sample-to-control relative abundance ratio of ≥ 5 and at least 20% prevalence in any single host species.

#### Distance calculations for multivariate regression on matrices

Multivariate regression on matrices (MRM) uses distances as both explanatory and response variables. As explanatory variables we used host phylogeny, diet, habitat (marine or terrestrial), ruminant digestion, and host species identity, all converted into distance matrices. The phylogenetic distances between samples were calculated from the host consensus tree from vertlife.org, with conspecific samples assigned a distance of 0. Dietary distances were calculated as Euclidean distances based on the proportions of animal, fruit, seed and other plant content in diet. Both distance matrices were centred and scaled to make the model coefficients comparable. For the categorical variables (host species, habitat, ruminant digestion), the distances were encoded in values of 0 (same) and 1 (different). For instance, the host species distance matrix had 0 for samples from the same host species, and 1 for samples from different host species.

As response variables we used three metrics of microbiome dissimilarity: Jaccard distances on non-normalised abundances, Euclidean distances on CLR-normalised abundances (Aitchison), and Euclidean distances on PhILR-normalised abundances. To test at different taxonomic ranks of microbiome composition (species to phylum), CLR- and PhILR-normalised abundances had to be re-calculated after agglomerating raw abundances to the desired taxonomic level. For PhILR normalisations, this required collapsing the microbial phylogeny to the required taxonomic rank, using functions from the ape R package[15]. All microbiome distances were then centred and scaled.

### Supplementary Figures

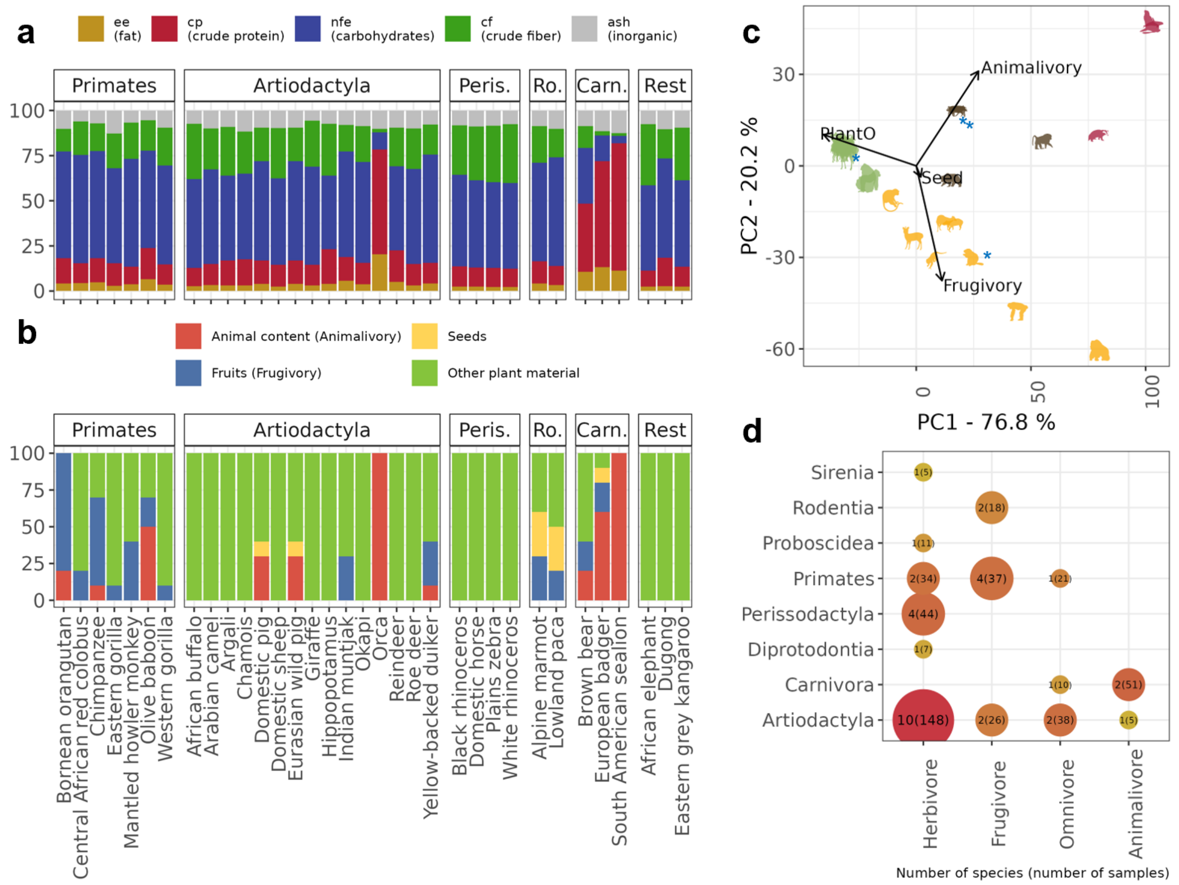

**Supplementary Fig. 1**. **Dietary diversity of the dataset. a**, Dietary composition of host species according to Lintulaakso et al.[16]. **b**, Dietary composition according to the EltonTraits database [17], after combining all dietary item types indicating animalivory (invertebrates, endotherm, ectotherm, unknown vertebrates and scavenge) into a single item type indicating animal material. Note that neither database included the domestic pig (*Sus domesticus*), so we used the same dietary data as for the European wild pig (*Sus scrofa*). We also used the older name for the Malayan Tapir*, Tapirus indicus*, instead of *Acrocodia indica* to match to the databases. **c,** PCA (Principal Components Analysis) ordination based on the data in panel b. PlantO = Other plant material. Animal figures were sourced using the rphylopic package[18] and coloured by dietary category. Blue asterisks indicate the four species with amended diet categories: the giraffe (frugivore to herbivore), the Eurasian wild pig and domestic pig (herbivore to omnivore), and the Alpine marmot (herbivore to frugivore). **d,** Distribution of host species (circle size, number outside brackets) and samples (circle colour, number in brackets) across taxonomic orders and dietary categories.

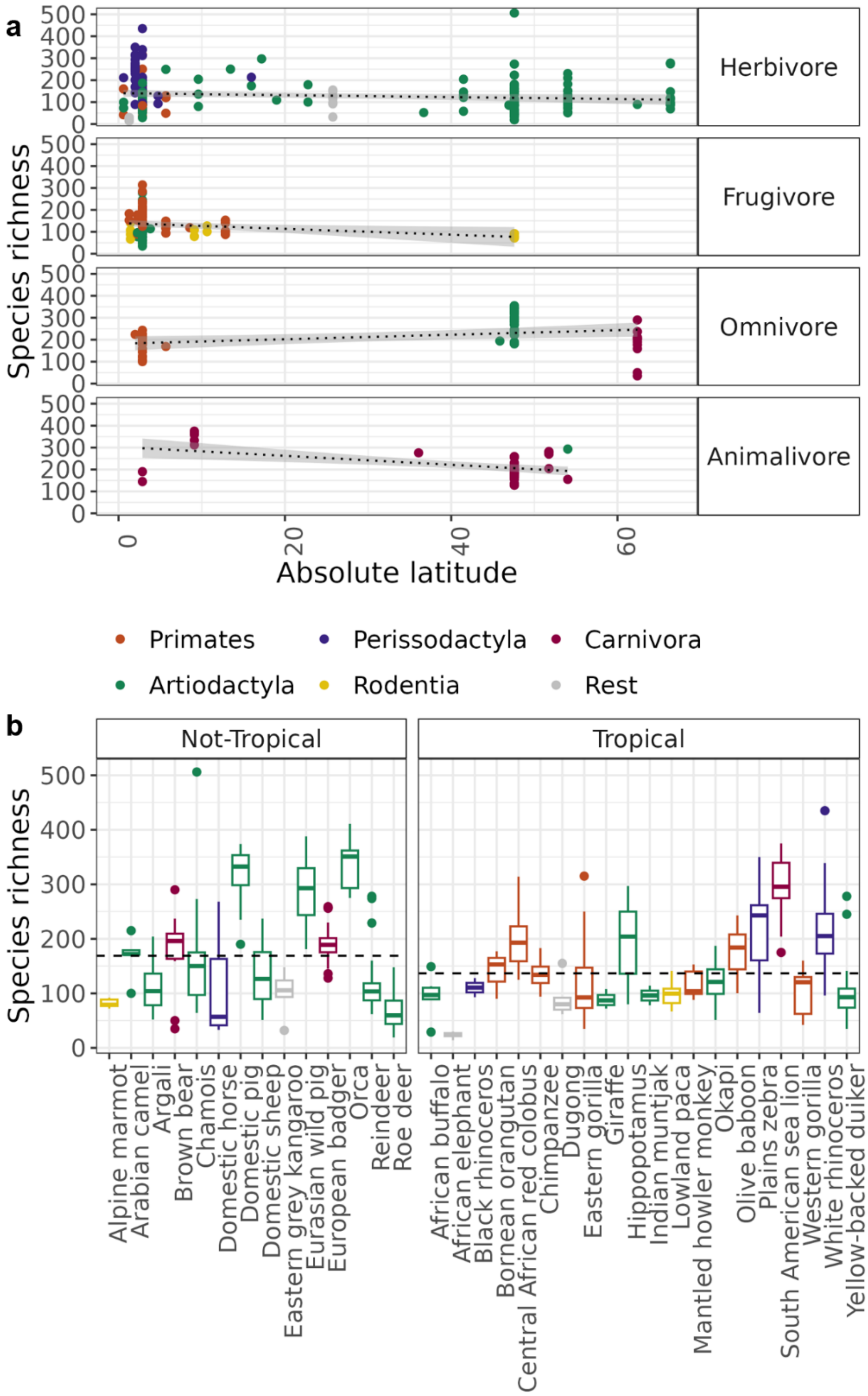

**Supplementary Fig. 2. No relationship between latitude and oral microbiome diversity. a**, Species richness plotted against the absolute value of estimated latitude (based on collection metadata, Supplementary Table 1). The dotted lines represent linear models fit separately for each dietary category. Across the entire dataset, ignoring dietary category, no effect of absolute latitude on species richness was observed (linear model p-value = 0.971). **b**, Species richness in host species with tropical and non-tropical distributions. In both panels, colour reflects host taxonomic order. The dashed lines reflect the weighted average richness for each distribution type (richness first averaged within species and then across species). Tropical and non-tropical species showed no significant differences in species richness (linear model p-value = 0.325).

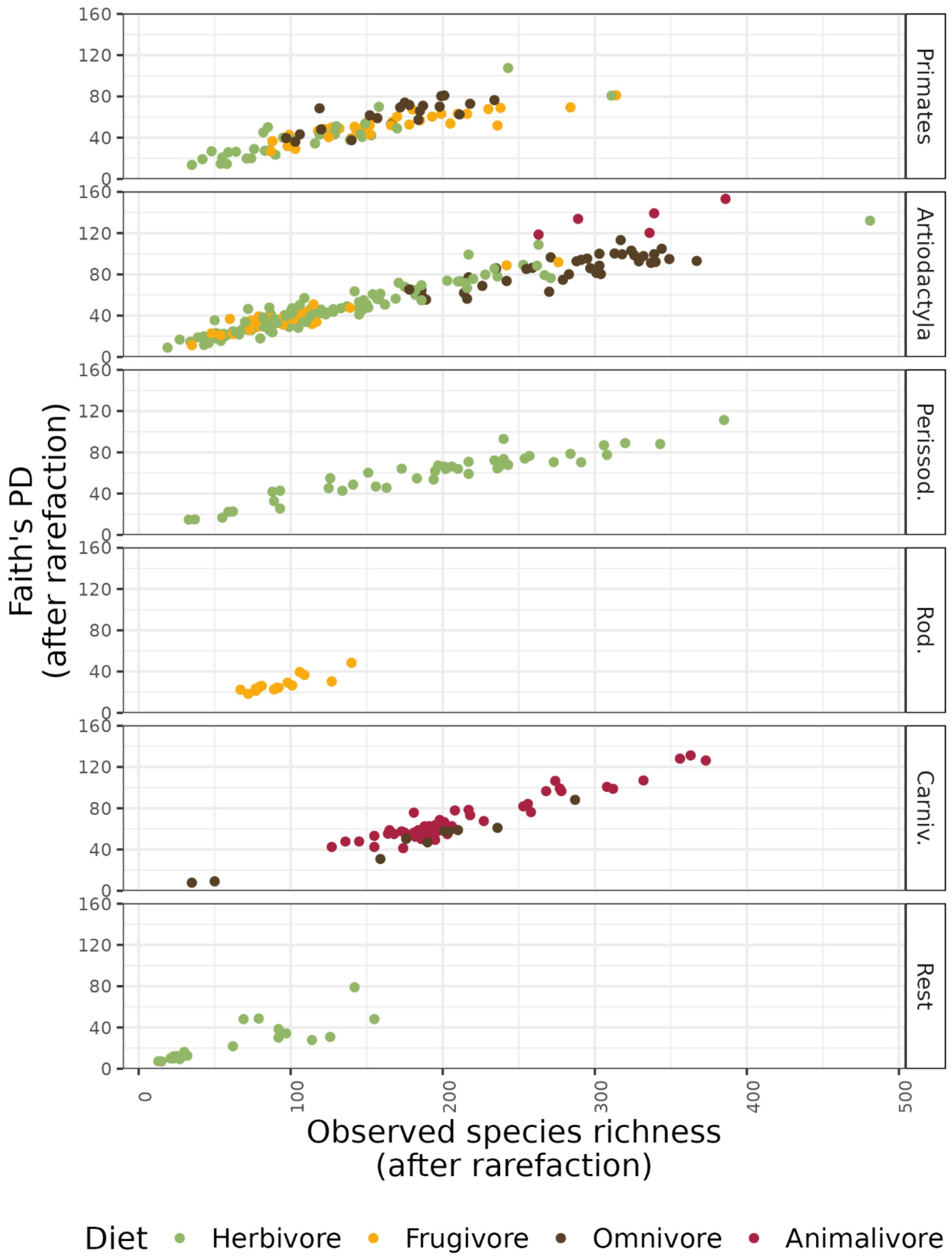

**Supplementary Fig. 3. Correlation between microbiome species richness and Faith's phylogenetic diversity.** Faith's PD plotted against observed species richness (r = 0.94, p-value < 0.001). Each point represents a sample and is coloured by dietary category (red = animalivore, brown = omnivore, yellow = frugivore, green = herbivore).

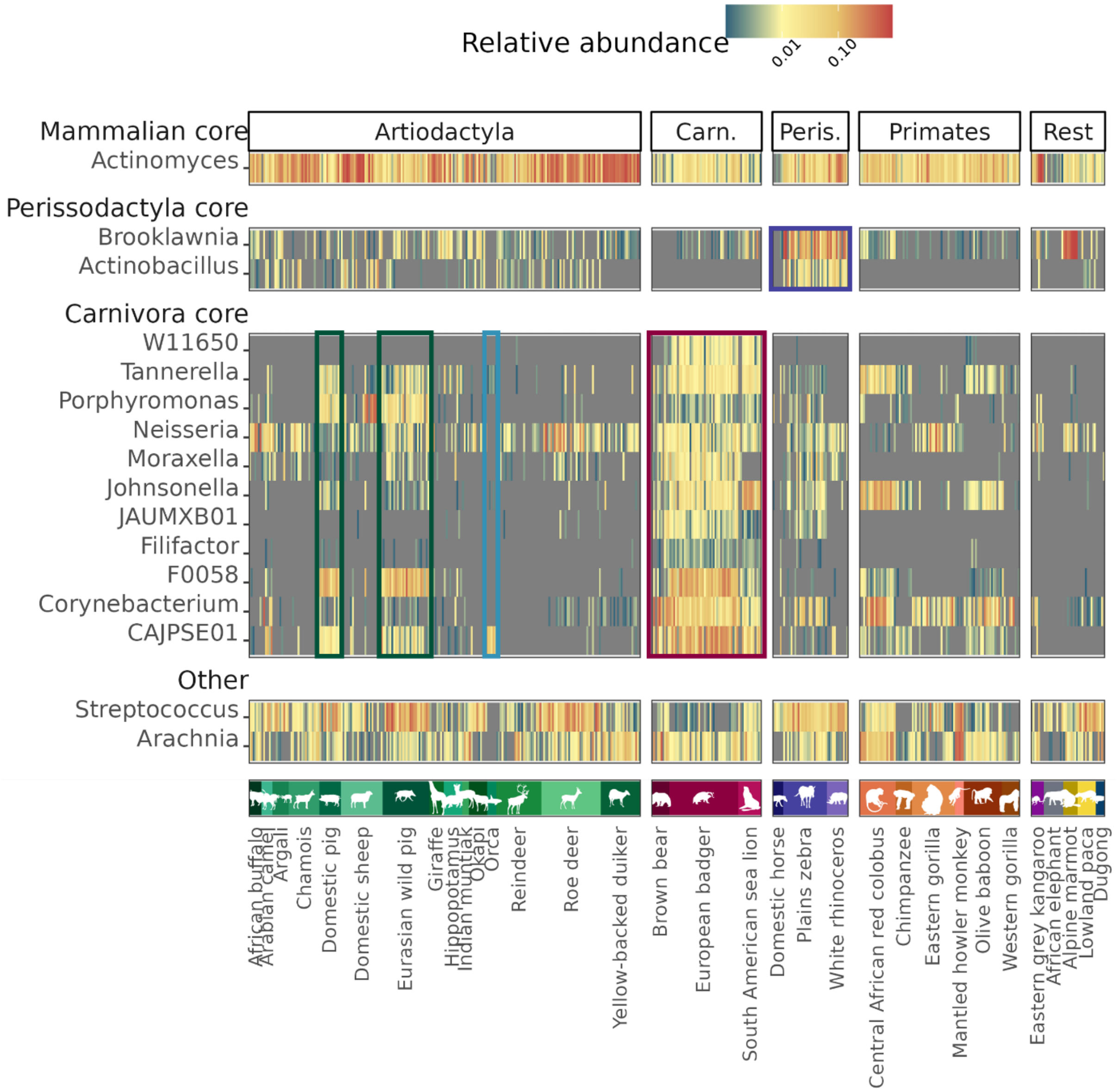

**Supplementary Fig. 4. Core genera across four mammalian orders with at least three species each.** Relative abundance of microbial genera that were identified as members of the core microbiome of one or more mammalian orders. Perissodactyla and Carnivora samples with their unique core microbiomes (excluding genera shared by other orders) are highlighted in purple and red, respectively. Note the greater abundance of Carnivora core taxa in omnivorous Artiodactyla (*Sus domesticus, Sus scrofa*), highlighted in green, compared to the animalivorous *Orcinus orca*, highlighted in blue. The colour scale is log-transformed and abundances below the detection threshold of 0.001% are shown in grey. Note that only host species with at least five samples are displayed here, excluding the black rhinoceros and the Bornean orangutan.

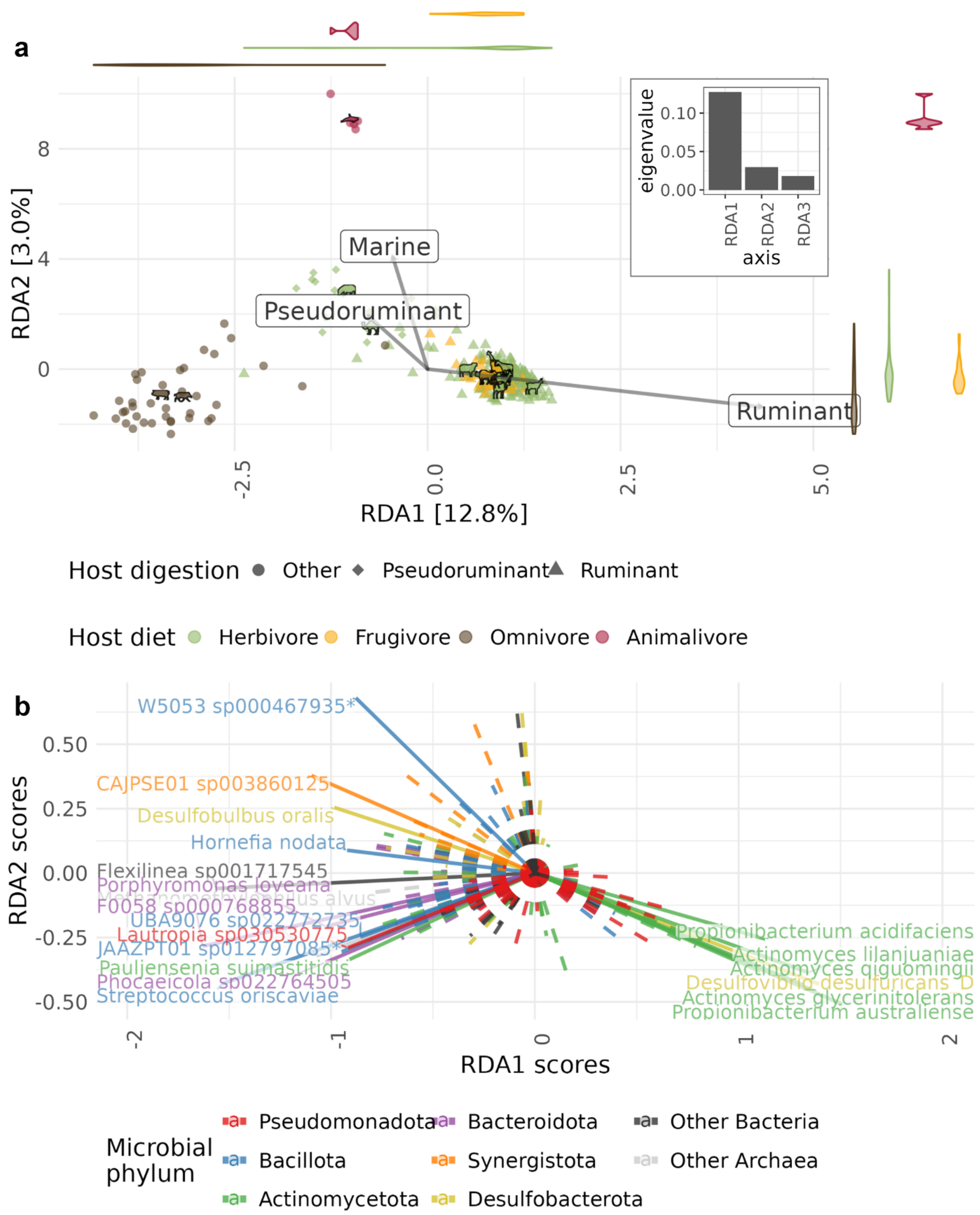

**Supplementary Fig. 5. Redundancy analysis on CLR-normalised abundances of species-level taxa for Artiodactyla samples. a**, Points represent samples and are coloured by dietary category, with shapes reflecting digestive physiology. The length of arrows indicates the effect of the explanatory variables used in the RDA: ruminant (most host species), pseudoruminant (Arabian camel and hippopotamus), other (orca, Eurasian wild pig and domestic pig) (Supplementary Table 2). Pseudoruminants appear at an intermediate position between ruminants and omnivores. The violin plots along the axes show the distribution of different dietary categories. The animal icons represent the centroids of each species. The inset is a scree plot, showing the variance explained per axis. **b,**

Loadings of 20 microbial species with the strongest correlation to the RDA axes in panel A (solid line), coloured by microbial phylum. Loadings for other taxa are represented with dashed lines.

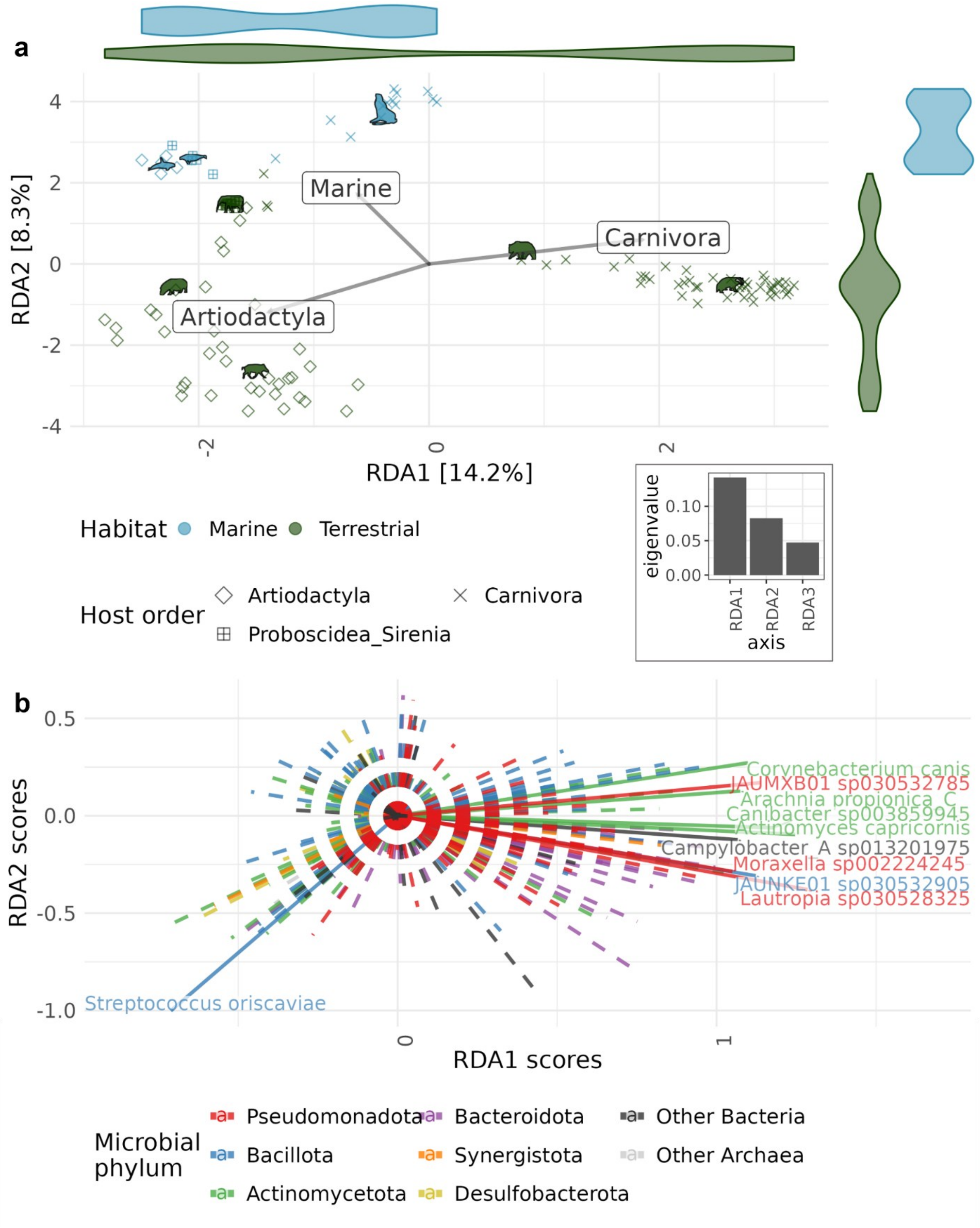

**Supplementary Fig. 6. Redundancy analysis on CLR-normalised abundances of species-level taxa for marine terrestrial comparisons. a**, The analysis was run for a subset of the data including marine mammals (orca, South American sealion, dugong) and their terrestrial relatives within our dataset (wild pig and hippopotamus, European badger, African elephant, respectively). Points represent samples and are coloured by marine or terrestrial habitat, with shapes reflecting host taxonomic order. Marine and terrestrial mammals are separated along the second axis, while Carnivorans are separated by the rest along the first axis. The length of arrows indicates the effect of the explanatory variables used in the RDA. The violin plots along the axes show the distribution of different dietary categories. The animal icons represent the centroids of each species. The inset is a scree plot, showing the variance explained per axis. **b,** Loadings of 15 microbial species with the strongest correlation to the RDA axes in panel A (solid line), coloured by microbial phylum. Loadings for other taxa are represented with dashed lines.

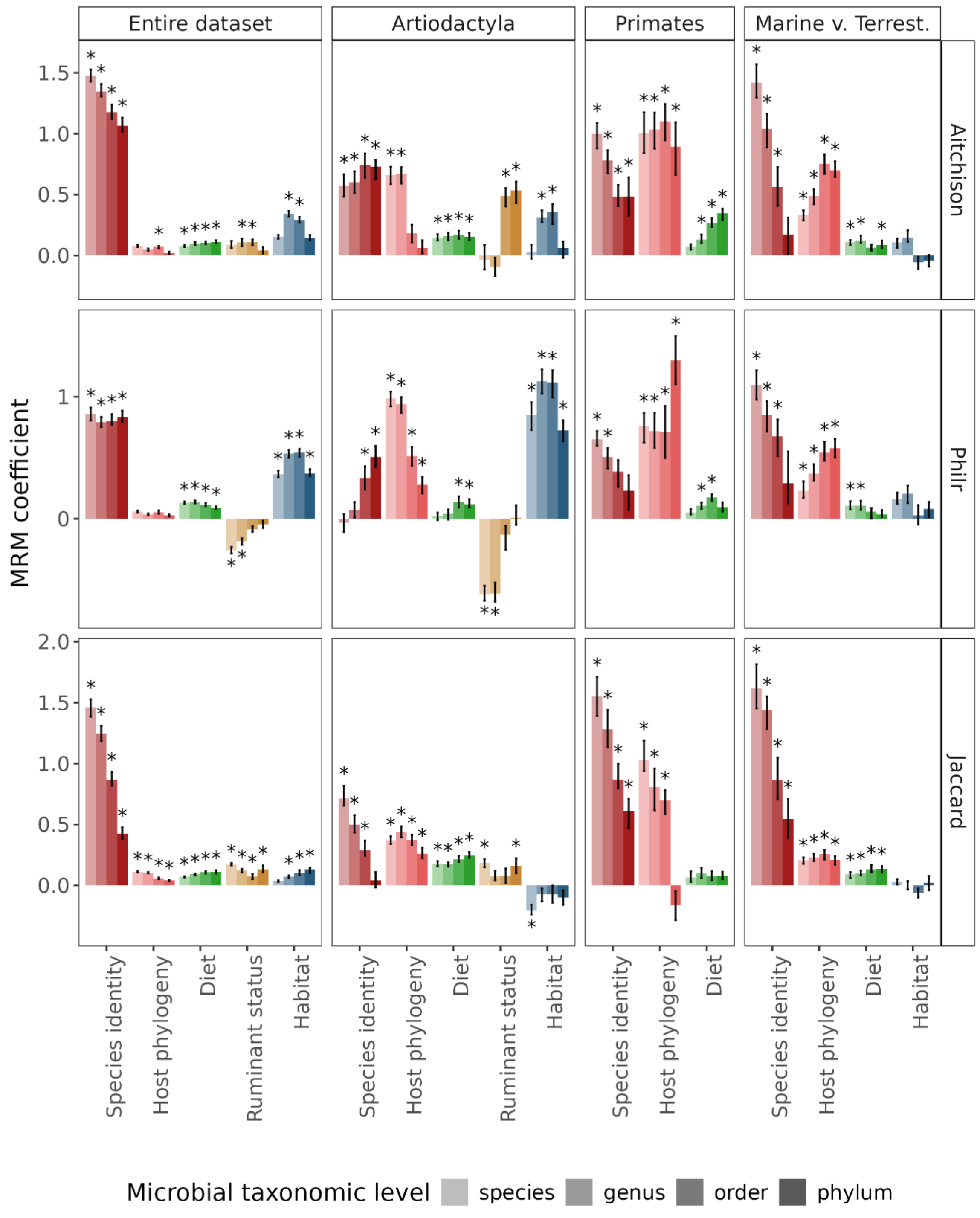

**Supplementary Fig. 7. Multiple regression on matrices shows host species identity as the main predictor of oral microbiome composition**. The analysis was run in the entire dataset and in subsets of Artiodactyla, Primates, and marine-terrestrial comparison sets. Three different microbiome distance metrics were used: Jaccard on non-normalised abundances, Aitchison (Euclidean on CLR-normalised abundances), and Euclidean on PhILR-normalised abundances, across four levels of microbial taxonomic resolution: species, genus, order, and phylum. The analysis was run iteratively 100 times for each model, randomly subsampling five samples per host species. The bar heights represent the median coefficient, and the error bars reflect the interquartile distance. The asterisks reflect a median p-value <0.05. In some cases, significantly negative results appear for habitat (marine or terrestrial) and ruminant status (ruminant or not), both of which are encoded as binarydistances (0 if two samples have the same value, and 1 if they do not). Therefore, these results suggest that two samples falling under the same category (e.g., both nonruminants) often have more different microbiomes than two samples that fall under different categories (e.g., a ruminant and a nonruminant).

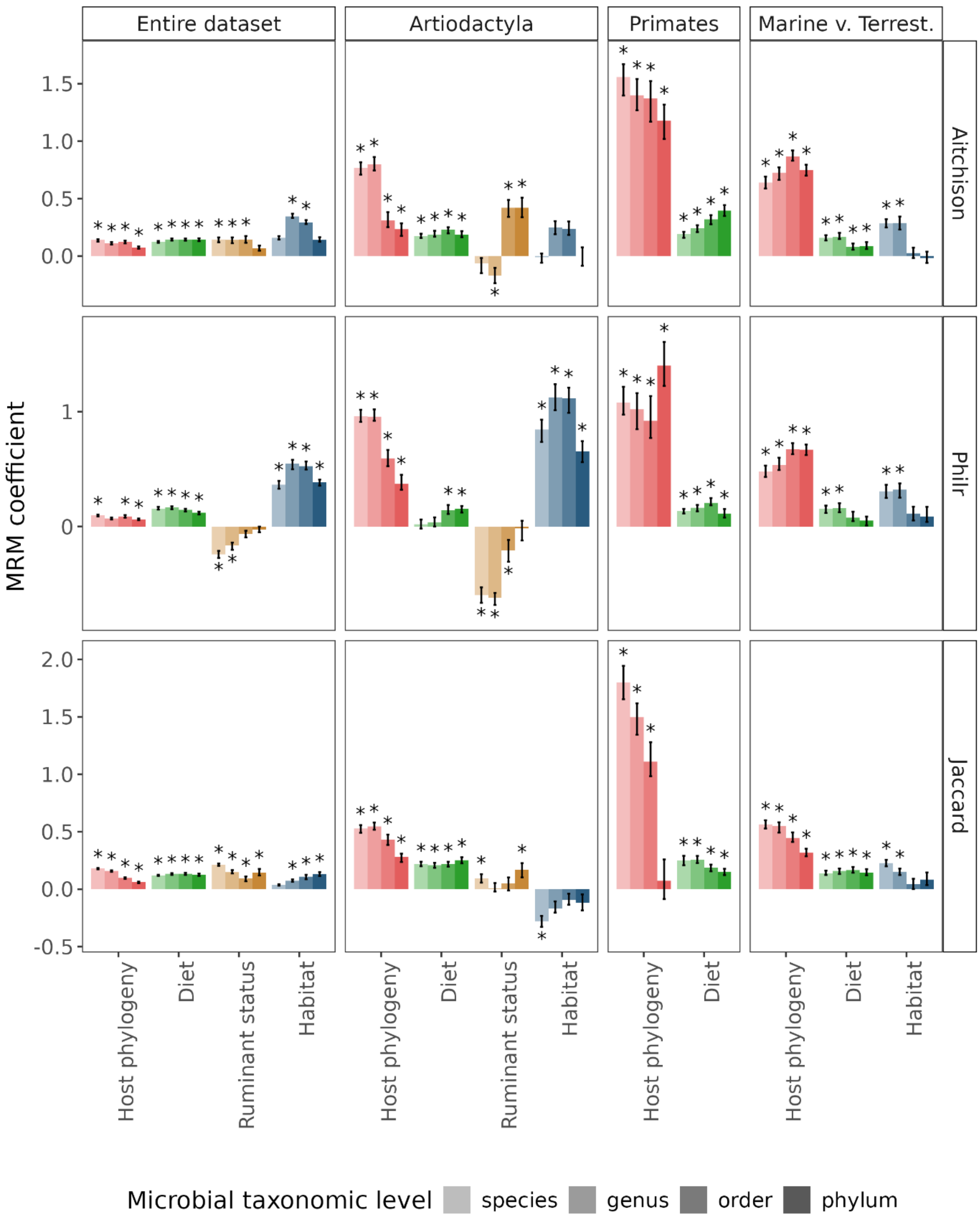

**Supplementary Fig. 8. Multiple regression on matrices without host species identity.** Removing host species identity as a predictor may lead to other predictors gaining statistical significance (e.g., host phylogeny in the entire dataset for Aitchison and Philr, habitat for the marine vs terrestrial subset). Overall, the relative contributions of the remaining predictors are consistent with Supplementary Fig. 7.

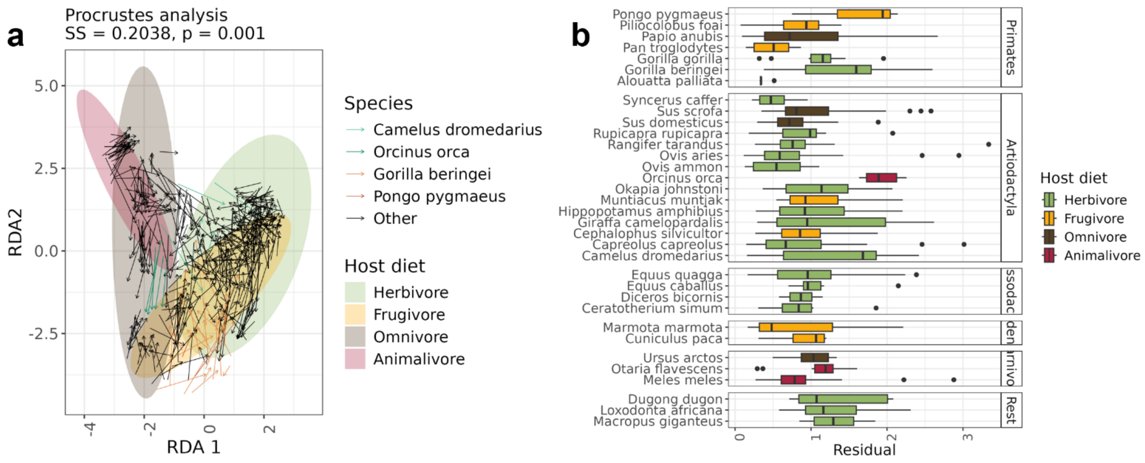

**Supplementary Fig. 9. Concordance between taxonomic and functional profiles of the oral microbiome. a**, Visualisation of Procrustes analysis comparing taxonomic (CLR-normalised genus abundances) and functional (CLR-normalised gene abundances) compositions. Each arrow represents a sample pointing from the position in the taxonomic RDA space to the position in the functional RDA space (rotated to maximise fit) and is coloured to highlight the host species with the largest residuals (from panel b). The ellipses represent dietary categories in the taxonomic ordination space. The two datasets show significant concordance according to the ‘protest’ statistical test (SS = 0.2038, p-value = 0.001). **b**, Residuals of the Procrustes analysis for each host species.

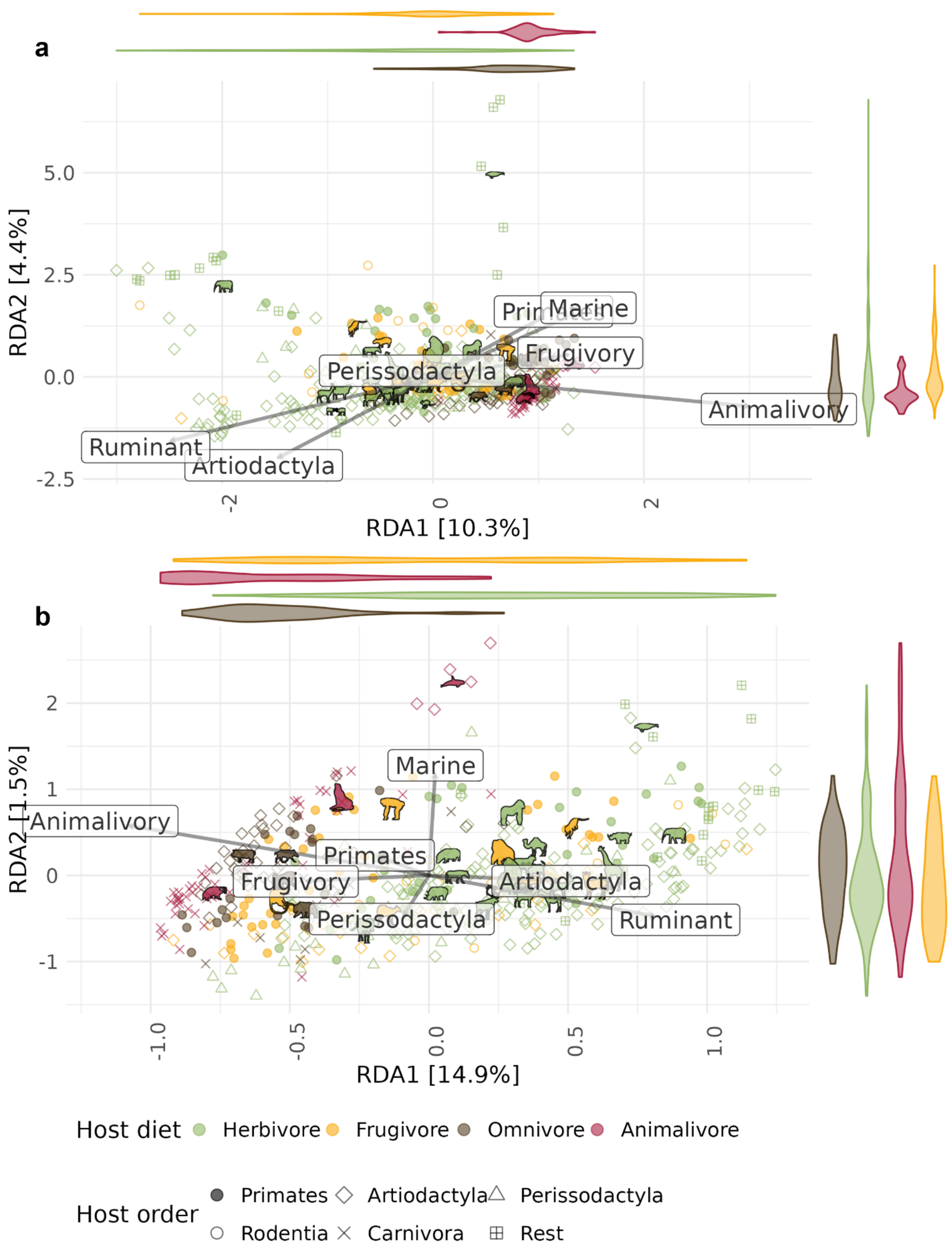

**Supplementary Fig. 10. Redundancy analysis on microbiome functional composition**. **a**, KEGG pathway abundances and b, functional trait completeness. Points represent samples, with colour signifying host dietary category and shape signifying taxonomic order. The arrows indicate the effect of the constraints used for the analysis, specifically continuous variables describing diet (animalivory, frugivory), marine habitat, ruminant digestion, and the most represented taxonomic orders (Artiodactyla, Perissodactyla, Primates). Violin plots alongside the axes show the distribution of different dietary categories.

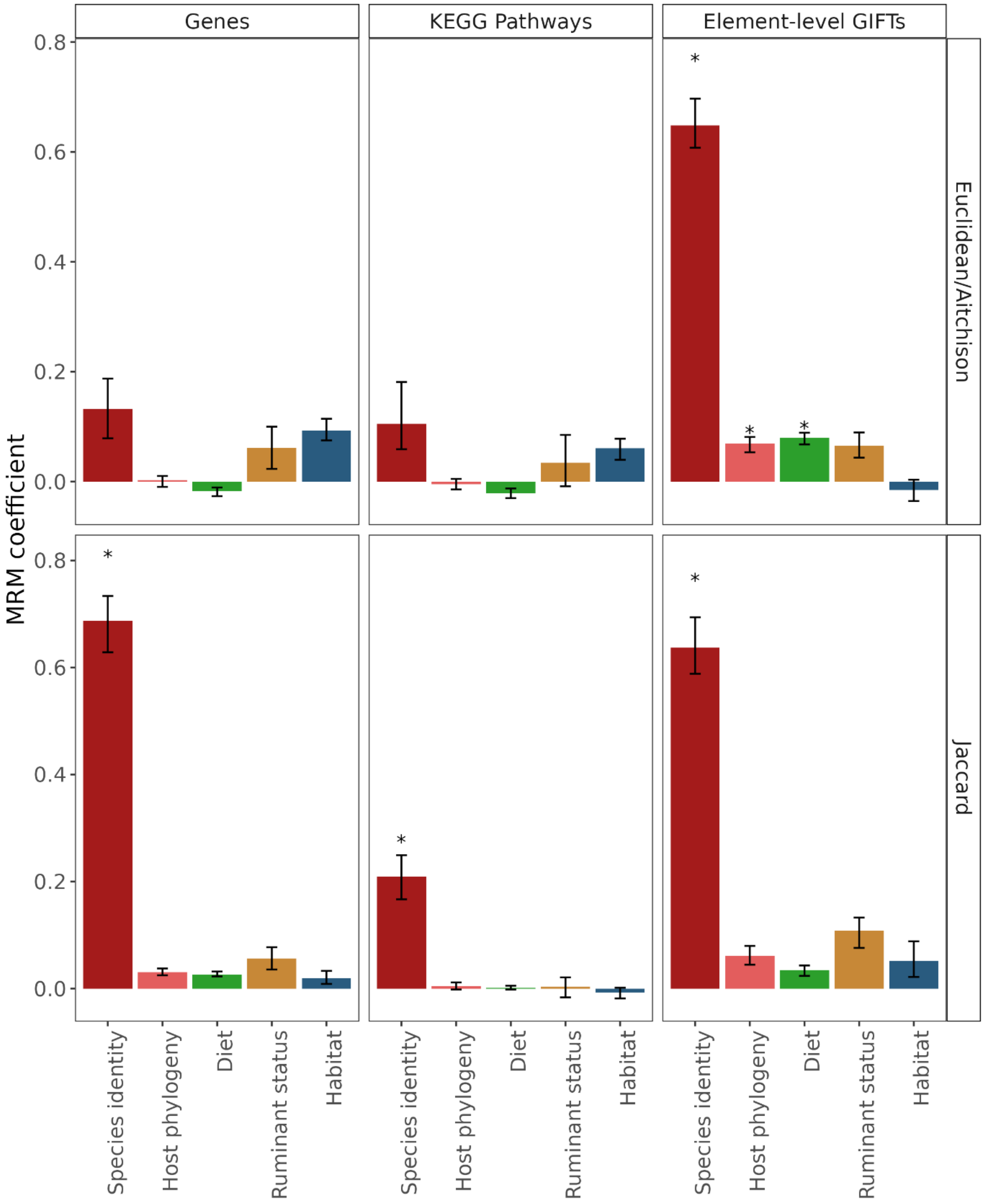

**Supplementary Fig. 11. Host species identity is the main predictor of functional composition.** Multiple regression on matrices (MRM) using abundances of genes, KEGG pathways, and completeness of genome-inferred functional traits at the element level. For pathways and genes, the analysis was run for both Jaccard and Aitchison distances. For functional traits, Euclidean distances were used instead of Aitchison, as their values did not reflect abundance, but completeness (values between 0 and 1). The analysis was run iteratively 100 times for each model, randomly subsampling 5 samples per host species. The bar heights represent the median coefficient and the error bars reflect the interquartile distance. The asterisks reflect a median p-value < 0.05.

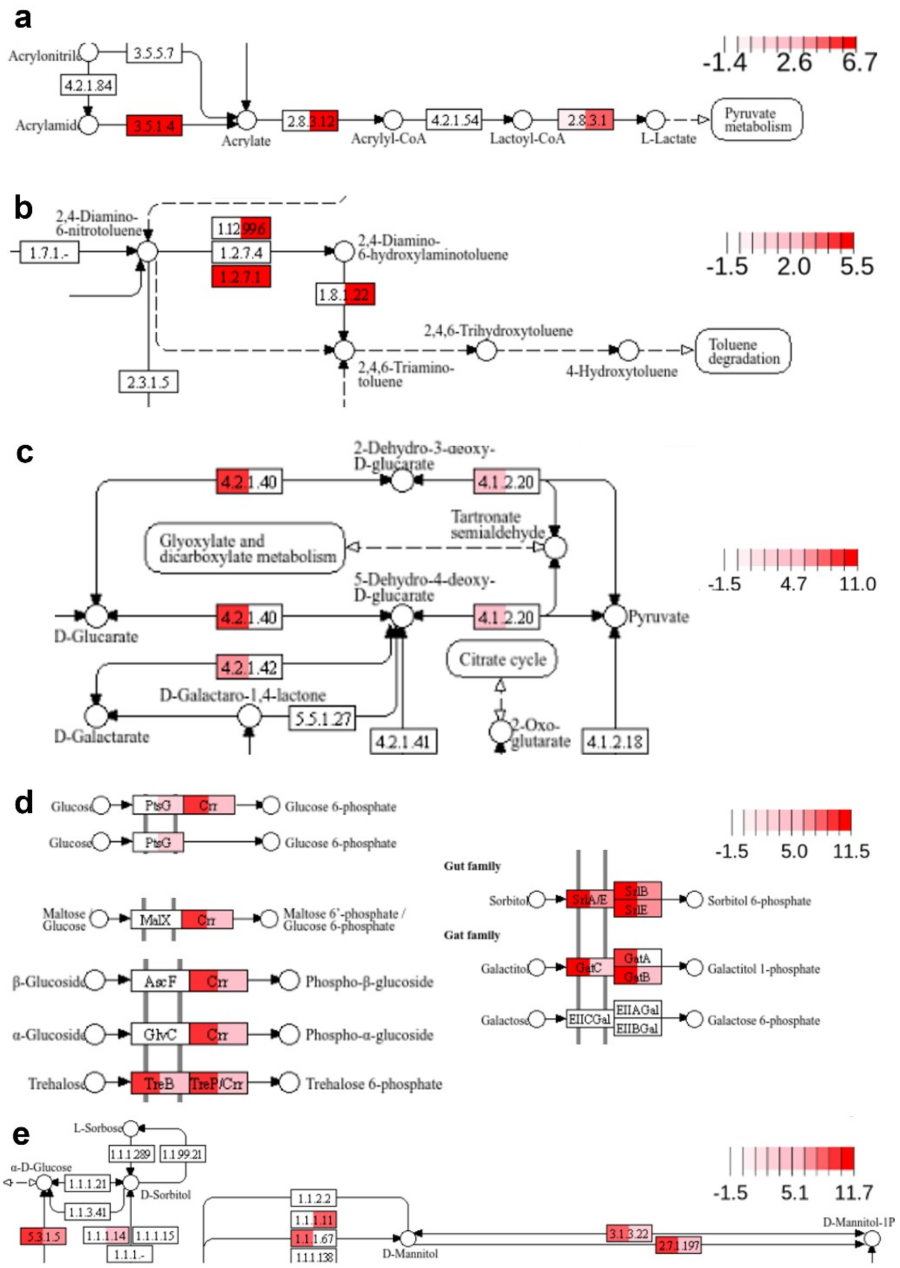

**Supplementary Fig. 12. Parts of KEGG pathways with high loadings on the pathway abundance RDA analysis.** Colour intensity shows the average abundance (CLR-normalised) of that gene in herbivores (left half) and carnivores (right half). Note different colour scales for each panel. **a**, Xenobiotic degradation: Styrene degradation (ko00643), highlighting the steps for acrylamid degradation, which were enriched in animalivores. **b**, Xenobiotic degradation: Nitrotoluene degradation (ko00633), highlighting steps enriched in animalivores. **c**, Ascorbate and aldarate metabolism (ko00053), highlighting glucarate/galactarate pathway, enriched in herbivores. **d,** Phosphotransferase system (PTS) (ko02060), highlighting sugar and sugar derivative transporters enriched in herbivores. **e,** Fructose and mannose metabolism (ko00051), highlighting mannitol metabolism steps, enriched in herbivores.

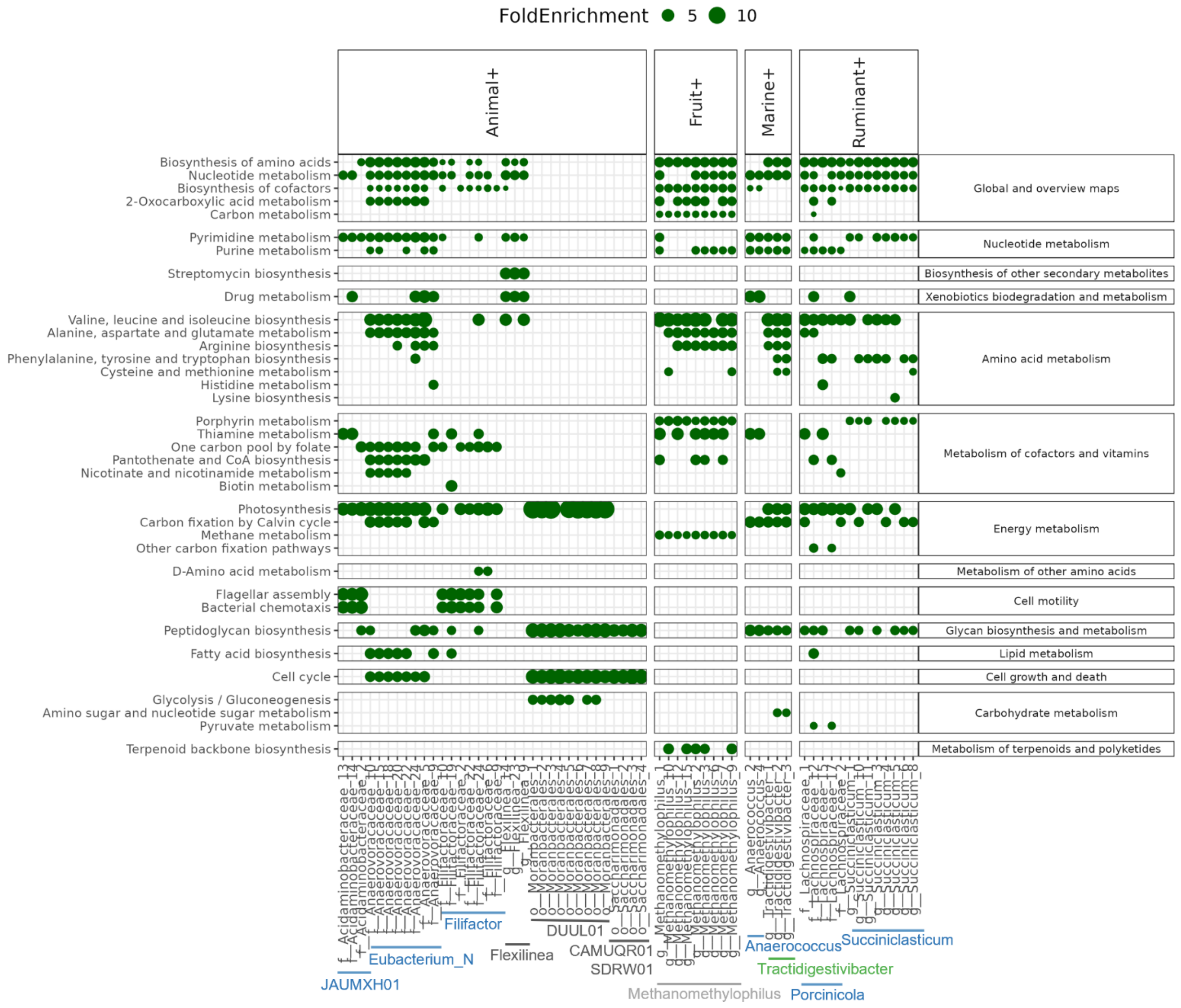

**Supplementary Fig. 13. Differential abundant taxa are enriched for functional pathways.** Gene set enrichment analysis for MAGs of taxa that showed increased abundance across the following factors: Proportion of animal and fruit in diet, marine lifestyle and ruminant digestion. Only categories "Metabolism" and "Cellular Processes" are shown. Note that the genomes assigned to the order Saccharimonadales could reflect two uncultured genera which are both enriched in animalivores, CAMUQR01 and SDRW01. The annotations are coloured based on the phylum a genus belongs to (blue: Bacillota, green: Actinomycetota, dark grey: other Bacteria, light grey: other Archaea).

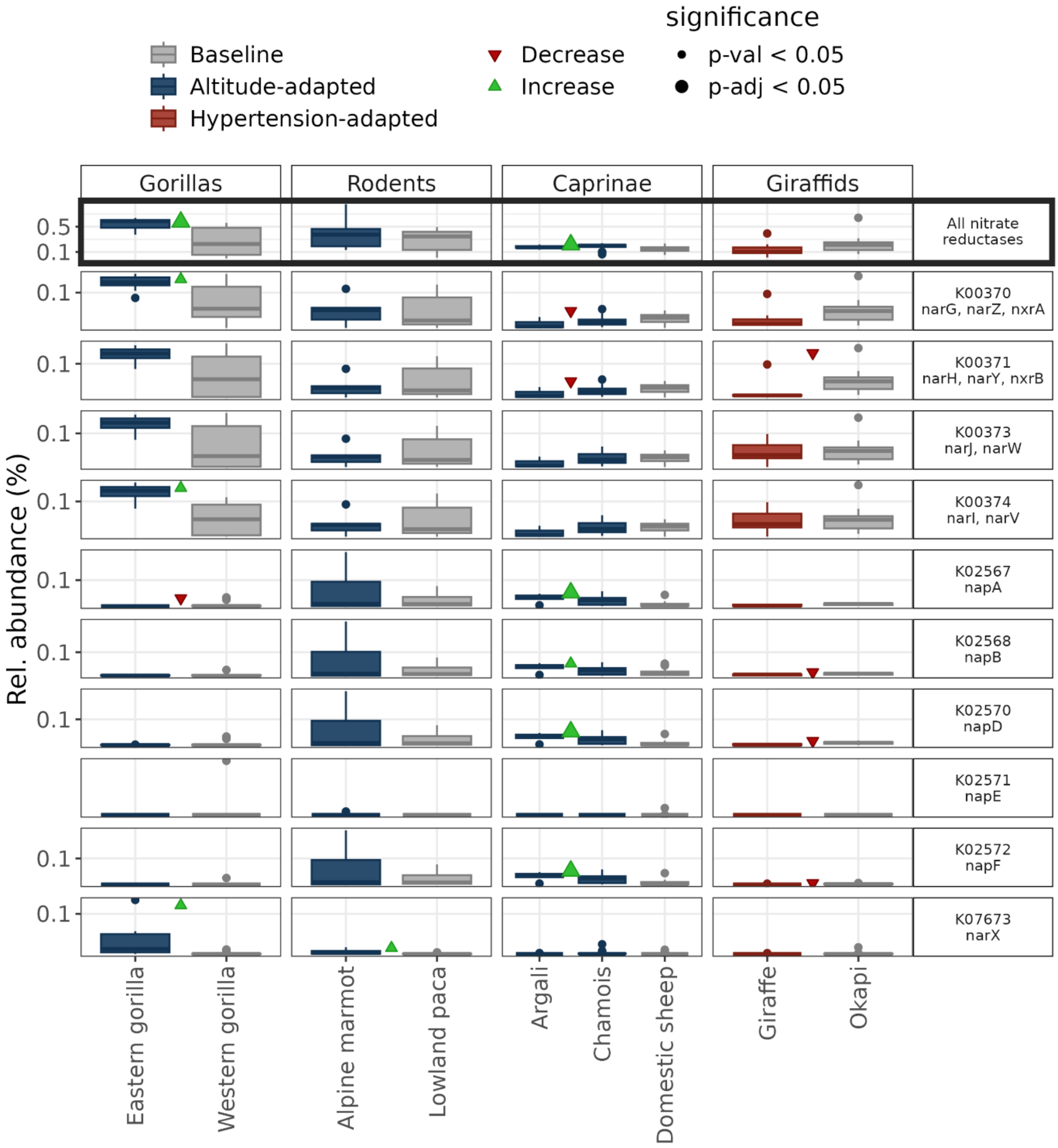

**Supplementary Fig. 14.** Relative abundances of genes encoding for nitrate reductases across three sets of host species, composed of altitude- or hypertension-adapted species (shown in dark blue or red, respectively), and a baseline species (in grey). The first row (highlighted) shows comparisons of the total relative abundance of relative reductase genes. The green arrowheads indicate a significant increase compared to the baseline species, according to a Wilcoxon test, and red arrowheads indicate a significant decrease. The subsequent columns show comparisons of individual nitrate reductase genes and were adjusted for multiple testing using the Holm method, with larger arrowheads indicating a p-value < 0.05 post-adjustment and the smaller ones indicating a p-value < 0.05 pre-adjustment. Argali and chamois were tested jointly against the domestic sheep. *nar*: membrane-bound respiratory nitrate reductase, *nap*: periplasmic nitrate reductase.

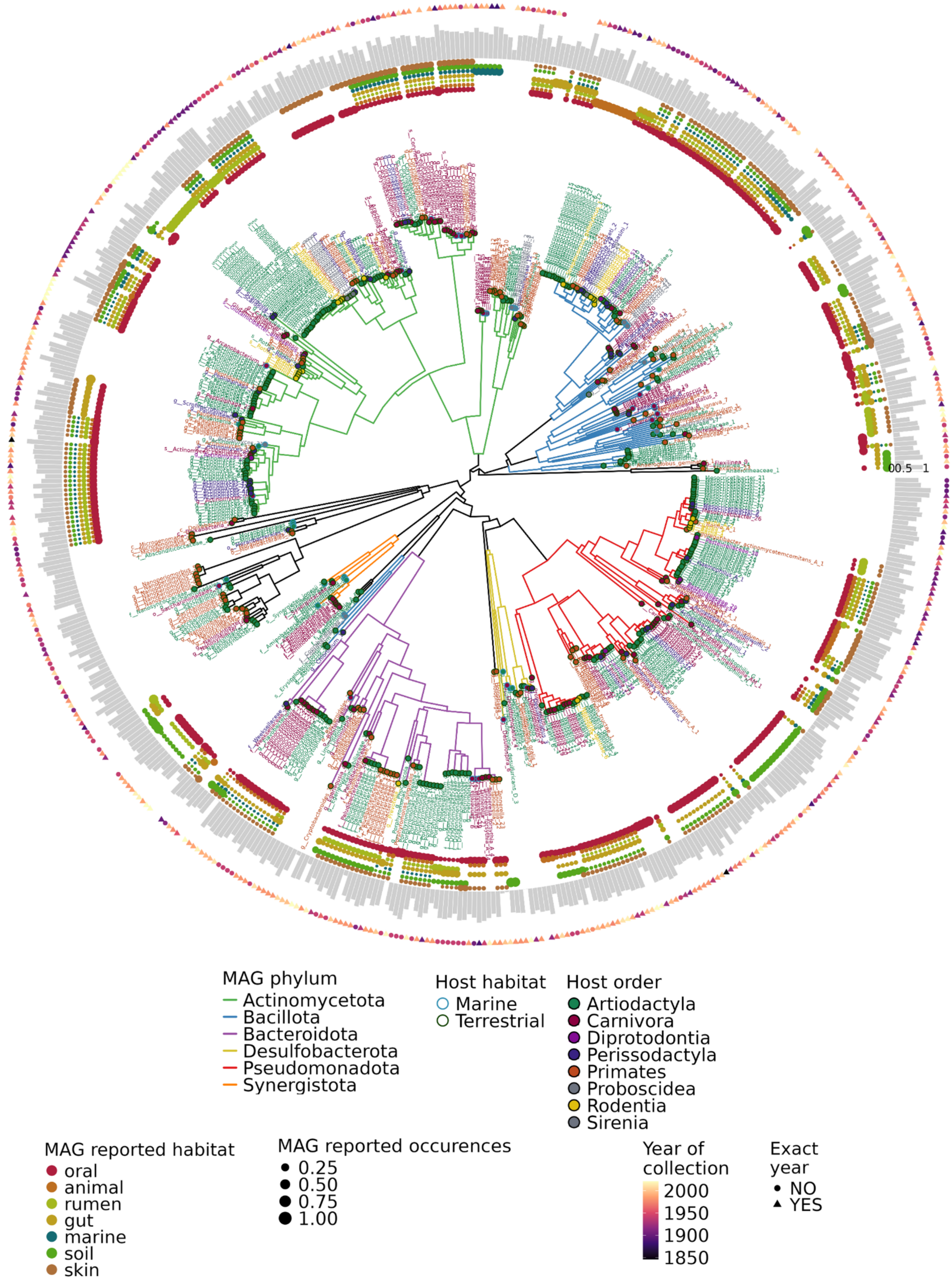

**Supplementary Fig. 15. Phylogenetic tree of 525 high-quality bacterial MAGs and 38 Patescibacteria MAGs.** High-quality MAGs had completeness ≥ 90% and contamination ≤ 5%, but Patescibacteria have reduced genomes, so they were retained regardless of completeness. From inside outwards: Tree branches are coloured by microbial phylum. Tree tip labels and tip circles are coloured according to the host taxonomic order from which the MAGs were assembled, with the circle outline denoting host habitat (marine or terrestrial). The dot plots represent different habitats that previously reported the bacterial taxa, according to the Omnicrobe database, starting from inner to outer circles with oral, animal, rumen, gut, marine, soil and skin. Circle size corresponds to the percentage of reports for the taxon in that habitat. The bars in grey represent the median amount of deamination damage at the 5’ end of the read (“damage_model_pmax” parameter from PyDamage). The outermost shapes are coloured according to the duration of preservation (age) of the specimen, with triangles if an exact collection year was available, and circles if the collection year had to be estimated.

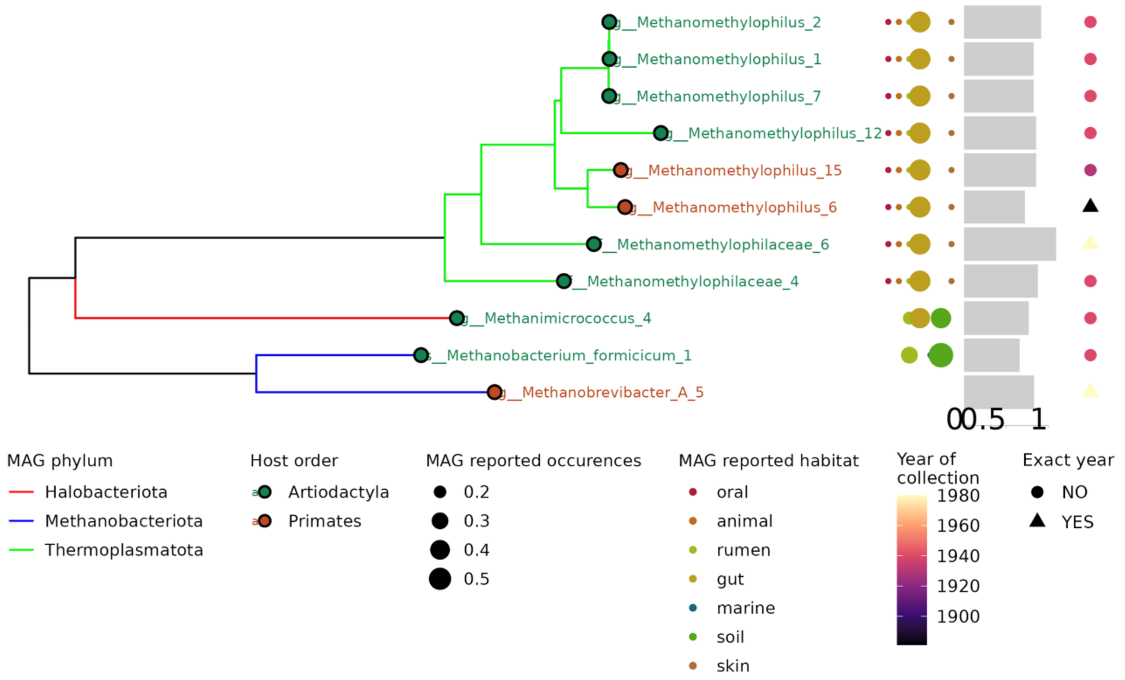

**Supplementary Fig. 16. Phylogenetic tree of 11 high-quality archaeal MAGs.** Phylogenetic tree of 11 high-quality archaeal MAGs (completeness ≥ 90% and contamination ≤ 5%). Tree branches are coloured by microbial phylum. The tree tip labels and tip circles are coloured according to the host taxonomic order from which the MAGs were assembled (Primates or Artiodactyla). From left to right: The dot plots represent different habitats that previously reported the archaeal taxon, according to the Omnicrobe database, starting from left to right with oral, animal, rumen, gut, marine, soil, and skin. Circle size corresponds to the percentage of reports for the taxon in that habitat. The bars in grey represent the median amount of deamination damage at the 5' end of the read ("damage_model_pmax" parameter from PyDamage). The outermost shapes are coloured according to the age of the specimen, with triangles if an exact collection year was available, and circles if the collection year had to be estimated.

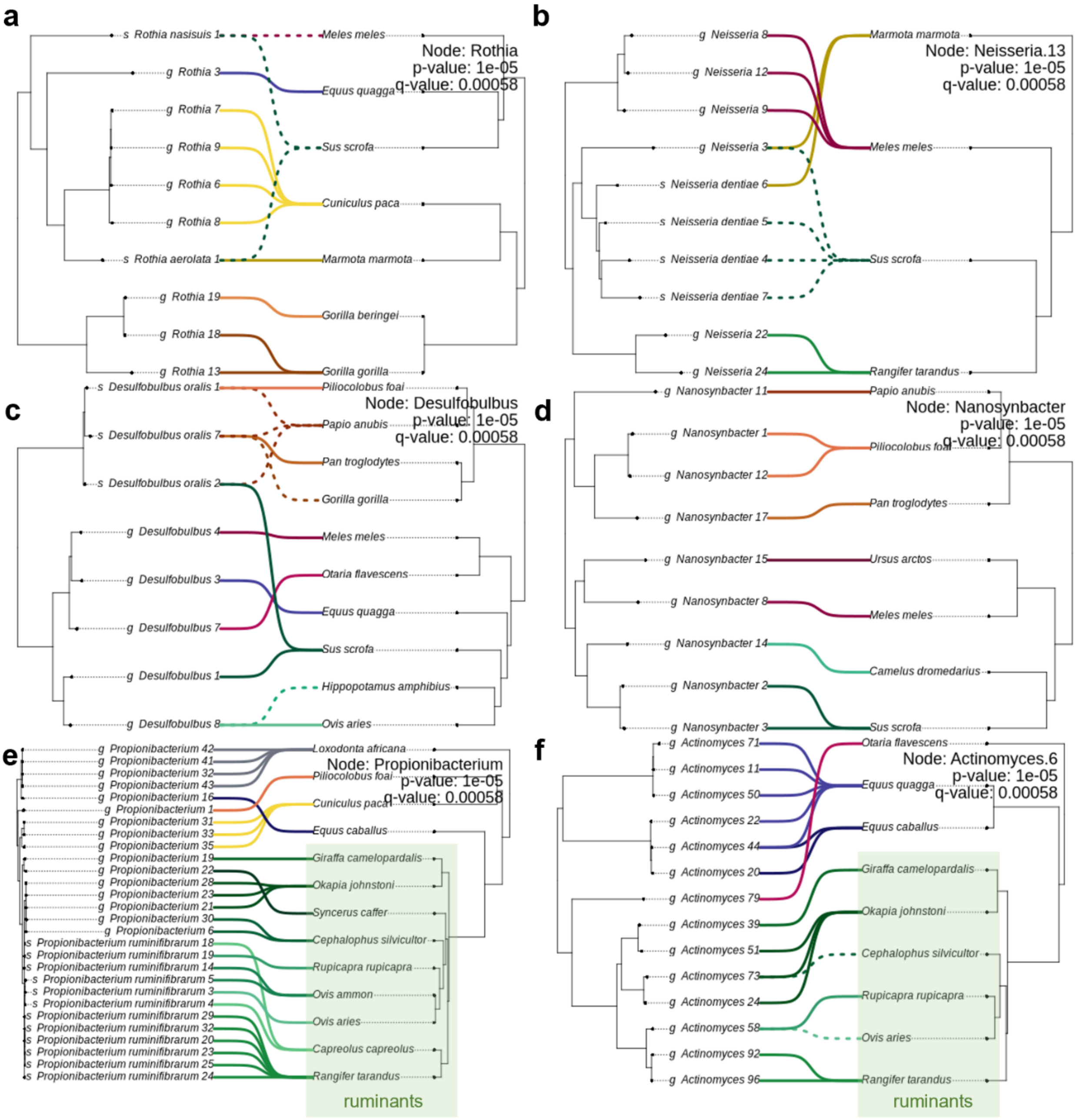

**Supplementary Fig. 17. Co-phylogenetic plots of select clades showing evidence of codiversification**. The MAG phylogeny is on the left and the host phylogeny on the right. Solid lines link MAGs to the host species they were assembled from, while dotted lines show MAG presence identified by mapping. In the top right corner, p-values before and after adjustment are displayed. **a**, *Rothia*; **b**, *Neisseria*.13; **c,** *Desulfobulbus*; **d**, *Nanosynbacter*; **e,** *Propionibacterium*; **f**, *Actinomyces*.6. In the last two panels, MAG clades assembled from ruminants are highlighted.

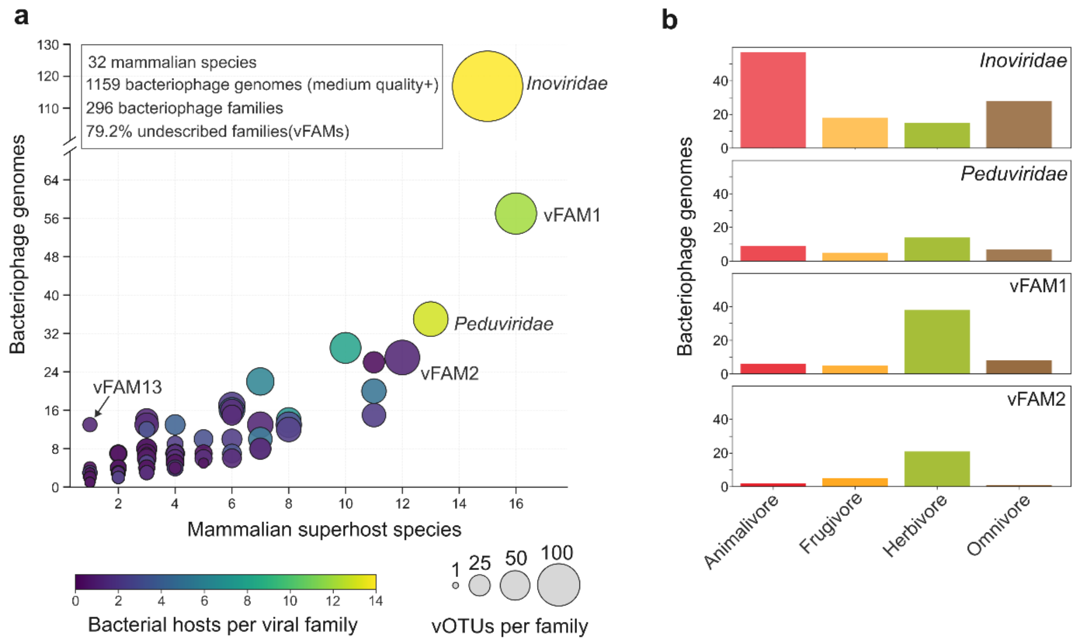

**Supplementary Fig. 18. Widespread presence of the dental calculus bacteriophage families in distantly related mammals. a**, Bubble plot showing the distribution of bacteriophage families across mammalian host species. Bubble size reflects the number of vOTUs per family and colour the number of distinct bacterial hosts. **b**, Distribution of the four largest viral families across mammalian host dietary categories. vFAM1 and vFAM2 were predominantly found in herbivores, whereas Inoviridae were enriched in animalivores.

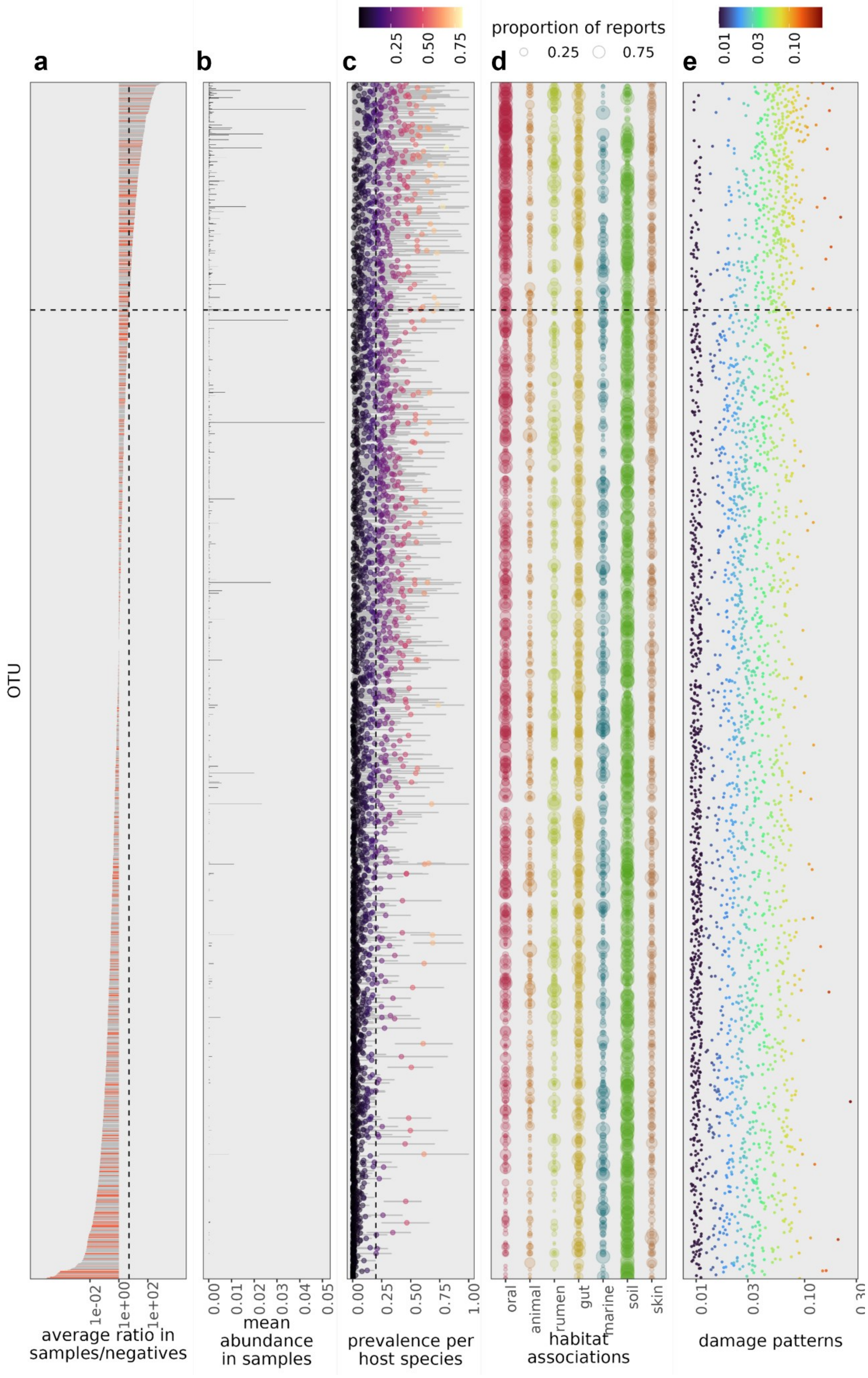
a
b
c
d
e
OTU
1e-02
1e+00
1e+02
average ratio in samples/negatives
0.00
0.01
0.02
0.03
0.04
0.05
mean abundance in samples
0.25
0.50
0.75
1.00
prevalence per host species
proportion of reports
oral
animal
rumen
gut
marine
soil
skin
habitat associations
0.01
0.03
0.10
damage patterns

**Supplementary Fig. 19**. **Identifying likely contaminants.** The y-axis shows all taxa found in both dental calculus samples and controls (N = 2687), ranked based on panel **a**, which shows the ratio of the (weighted) average relative abundance in samples to the average relative abundance in negative controls (including environmental samples and extraction and library preparation negatives), so that the taxa that are on average more abundant in samples than controls appear on top in all panels (note log10-transformed x-axis). The red bars indicate that the taxon belongs to a microbial genus previously reported as a common laboratory contaminant [9,19]. These laboratory contaminant lists are collated at the genus level, and genera may contain both oral and environmental taxa. The vertical dashed line signifies the ratio threshold of 5 that was used for decontamination. **b,** Weighted average abundance of each taxon in dental calculus samples (first averaged within host species, and then across host species, to account for differences in sample sizes). **c,** Prevalence of taxa in each host species (the proportion of samples with that taxon present at more than 0.01% relative abundance). Points indicate the average prevalence across all host species, and are coloured accordingly, and the grey bars show the minimum and maximum values. The vertical dashed line shows the 20% minimum prevalence threshold used for filtering. **d,** Previously reported habitats for each taxon from the Omnicrobe database. Pre-defined habitats were further grouped for easier visualisation: The oral habitat (red) includes the Omnicrobe habitat terms “dental plaque” and “mouth”; The animal habitat (orange) includes terms "mammalian", "wild animal" and "mammalian livestock"; rumen (light green) includes only the term “rumen”; gut (yellow) includes only the term “gut”; marine (blue) includes the terms "marine water" and "deep sea"; soil (green) includes only the term “soil”; skin (beige) includes only the term “skin”. Circle size corresponds to the percentage of reports for that habitat. **e,** Amount of deamination damage at the 5’ end of the read (“damage_model_pmax” parameter from PyDamage; note log10-transformed x-axis). Points indicate the median value for that taxon, and are coloured accordingly. Across all panels, the horizontal dashed line shows the cut-off used for decontamination, only retaining taxa above the line for downstream analysis. Retained taxa showed high sample prevalence, were often reported in oral environments and produced consistent post-mortem DNA damage patterns.

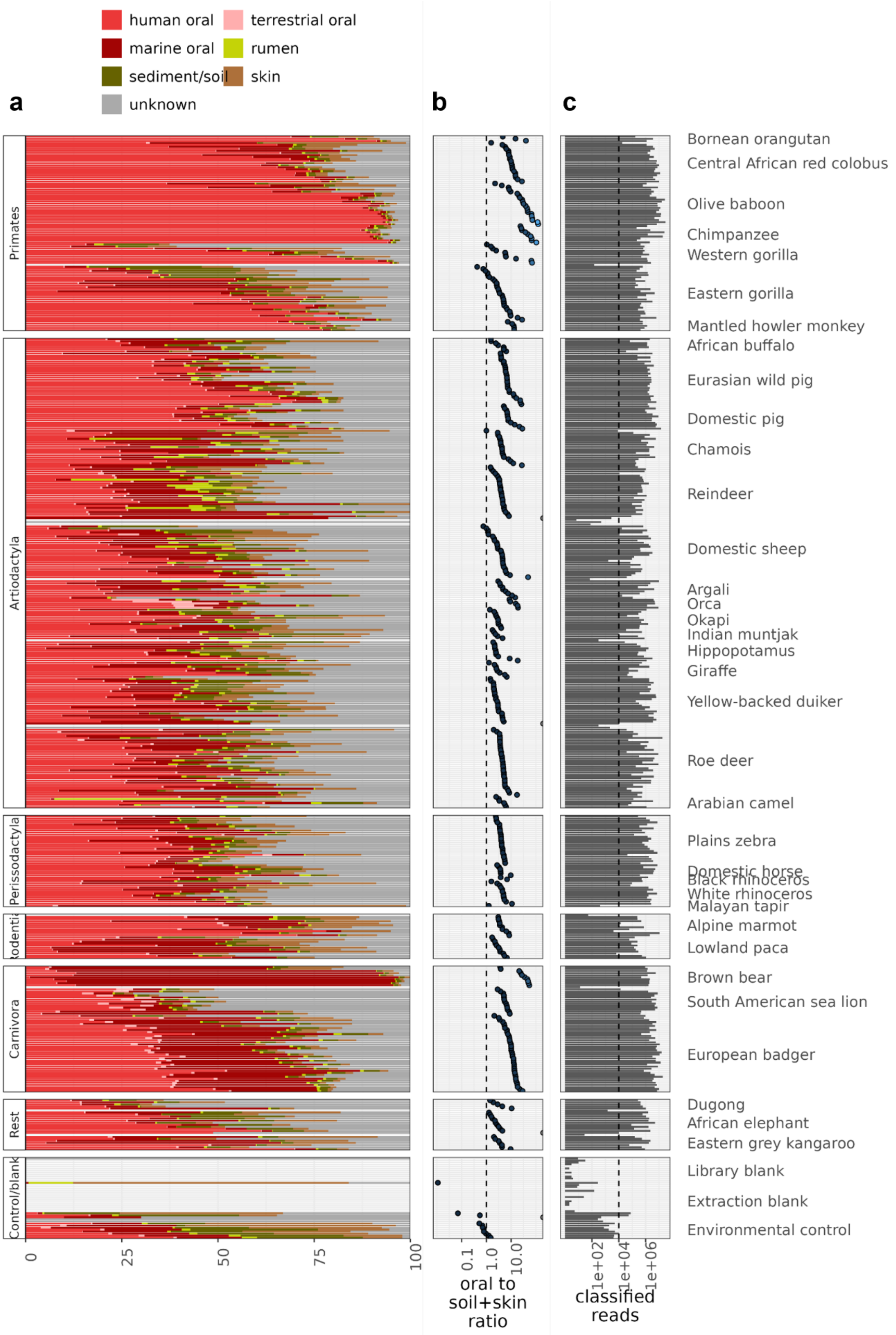

**Supplementary Fig. 20. Identifying poorly preserved samples. a,** Source tracking results obtained using decOM with a custom source matrix. Note some samples failed to be analysed by this tool and are shown in white. For these, only the read count filter was applied. **b,** The ratio of the oral microbiome proportion (combining all oral fractions) to the proportion of likely contaminants (skin and soil microbiomes) as seen in the first panel. Samples with ratio to the left of the parity line were removed from further analyses. **c,** The number of classified reads in each sample with the dashed line corresponding to the cut-off of 10,000 classified reads for retained samples.

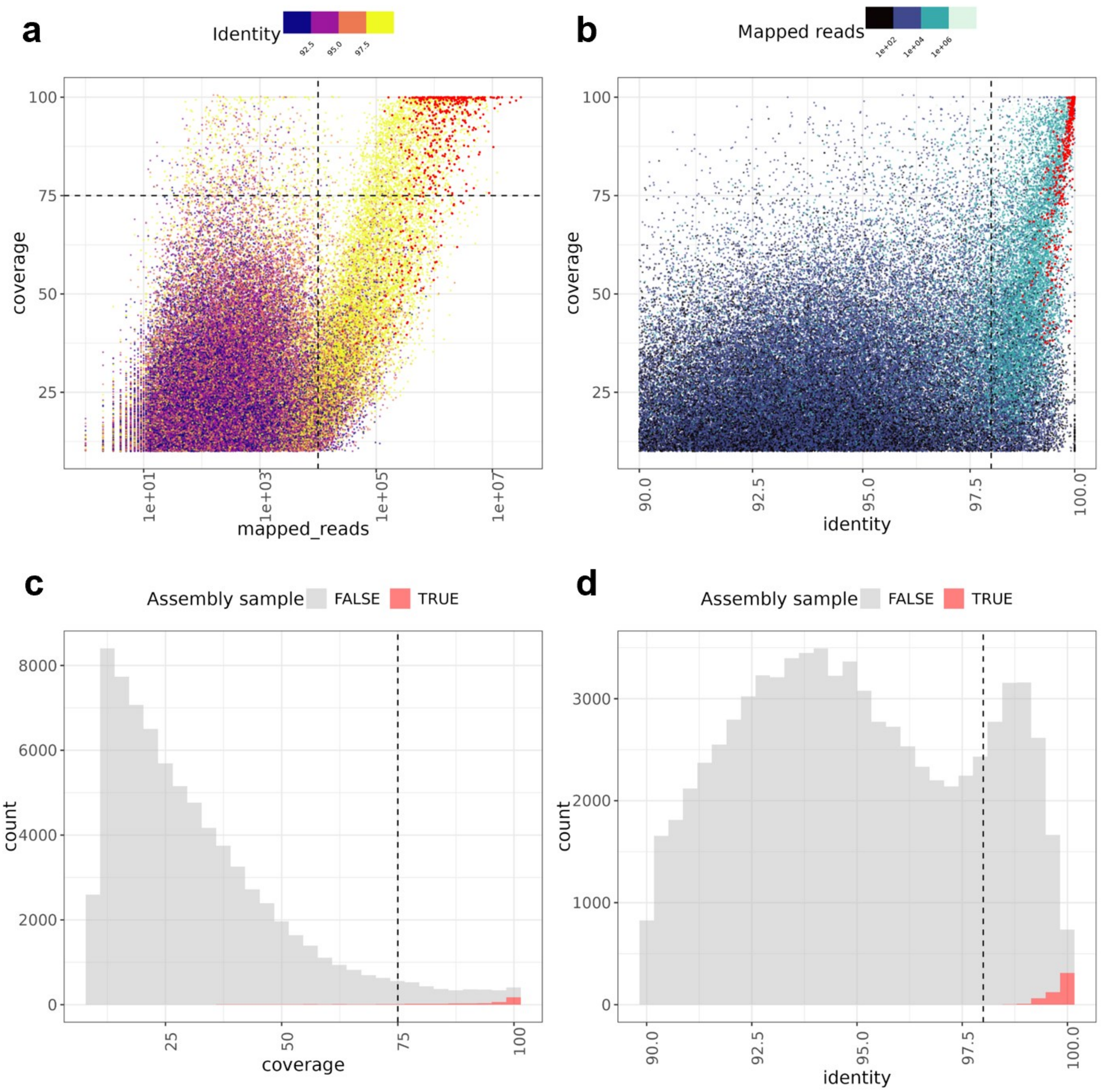

**Supplementary Fig. 21. Evaluating MAG presence in host samples.** Every data point represents the mapping of one metagenome to one high quality MAG (completeness ≥ 90% and contamination ≤ 5%). Only mappings with at least 90% average query nucleotide identity (the percentage of nucleotides in the mapped reads that are identical to the MAG) are displayed. **a,** Breadth of coverage (percent of the MAG sequenced that is covered by at least one read) against the total of number of mapped reads, coloured by average query nucleotide identity. The vertical and horizontal dashed lines represent the used cut-offs for number of mapped reads and breadth of MAG coverage, respectively. **b,** Breadth of coverage plotted against average query nucleotide identity, coloured by number of mapped reads. The dashed vertical line represents the 98% nucleotide identity threshold used to consider a MAG present in that sample. In both **a** and **b**, the red points represent within-metagenome mappings (reads from a metagenome being mapped to a MAG which was assembled from that same metagenome). **c,** Distribution of breadth of coverage values for all mappings. **d,** Distribution of average query nucleotide identity for all mappings. Within-metagenome mappings represent in red, whereas cross-metagenome mappings are shown grey.

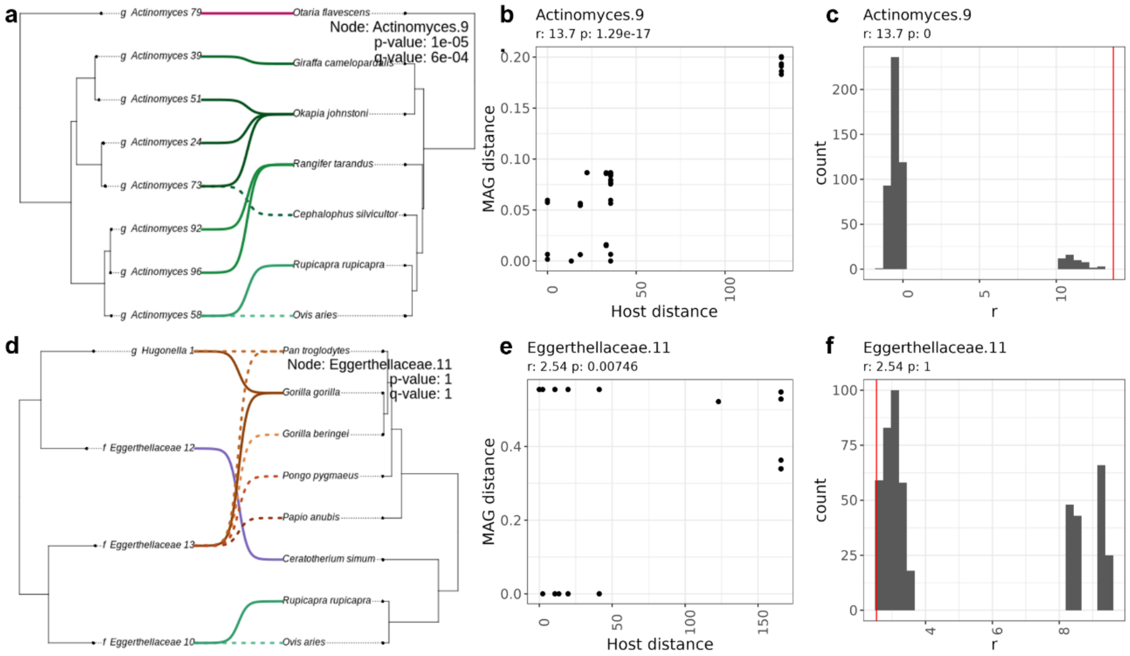

**Supplementary Fig. 22. Testing for co-diversification between MAGs and host species.** Top row: Actinomyces.9, a clade showing evidence for co-diversification. Bottom row: Eggerthellaceae.11, a clade without evidence of co-diversification. **a,** and **d,** Co-phylogenetic plots, with the MAG phylogeny on the left and the host phylogeny on the right. Solid lines link MAGs to the host species they were assembled from, whereas dotted lines show MAG presence identified by mapping. P-values (before and after adjustment with the Holm method) based on the permutation test are also displayed. **b,** and **e,** MAG phylogenetic distances plotted against host phylogenetic distances, with p-value from a Pearson correlation test. **c,** and **f,** Distributions of the correlation coefficient r, after permuting the MAG tree tips 500 times. The red line indicates the value of r for the original unpermuted tree. The p-value displayed here reflect the proportion of samples with r values equal to or greater than the original r.

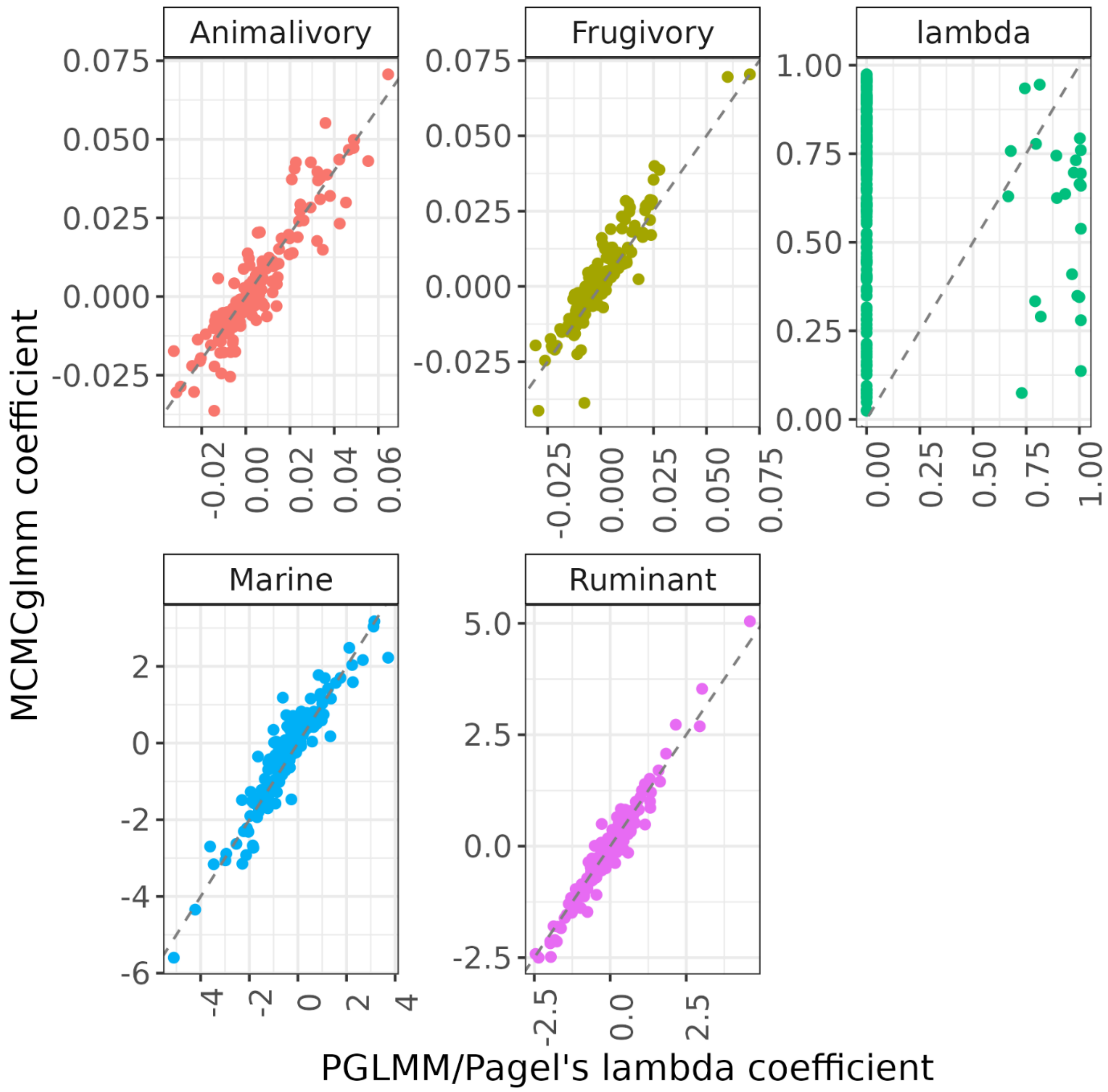

**Supplementary Figure 23. Correlation between coefficient estimated by PGLMM and MCMCglmm.** The estimated coefficients from traditional phylogenetic methods (PGLMM for the fixed effect Animal, Fruit, Marine and Ruminant, as well as Pagel's lambda for the effect of phylogeny) and the Bayesian approach MCMCglmm. Every point represents one of the 200 microbial genera used in these analyses. The estimates of the fixed effects are strongly correlated, but not those of the phylogenetic effects.
